# Overcoming immune evasion from post-translational modification of a mutant KRAS epitope to achieve TCR-T cell-mediated antitumor activity

**DOI:** 10.1101/2024.09.18.612965

**Authors:** Jihoon William Lee, Emily Y. Chen, Taylor Hu, Rachel Perret, Mary E. Chaffee, Tijana Martinov, Shwetha Mureli, Clara L. McCurdy, Lisa A. Jones, Philip R. Gafken, Pritha Chanana, Yapeng Su, Aude G. Chapuis, Philip Bradley, Thomas M. Schmitt, Philip D. Greenberg

**Affiliations:** Program in Immunology, Division of Translational Science and Therapeutics, Fred Hutchinson Cancer Center, Seattle, WA, USA; Medical Scientist Training Program, University of Washington, Seattle, WA, USA; Proteomics and Metabolomics Shared Resource, Fred Hutchinson Cancer Center, Seattle, WA, USA; Genomics and Bioinformatics Shared Resource, Fred Hutchinson Cancer Center, Seattle, WA, USA; Herbold Computational Biology Program, Fred Hutchinson Cancer Center, Seattle, WA; Department of Medicine, Division of Hematology and Oncology, University of Washington, Seattle, Washington, USA; Department of Immunology, University of Washington, Seattle, WA

**Author notes:** Corresponding author(s): Philip Greenberg, MD, Thomas Schmitt, PhD. Co-senior authors who contributed equally (Thomas M. Schmitt, Philip D. Greenberg).

## Abstract

T cell receptor (TCR)-T cell immunotherapy, in which T cells are engineered to express a TCR targeting a tumor epitope, is a form of adoptive cell therapy (ACT) that has exhibited promise against various tumor types. Mutants of oncoprotein KRAS, particularly at glycine-12 (G12), are frequent drivers of tumorigenicity, making them attractive targets for TCR-T cell therapy. However, class I-restricted TCRs specifically targeting G12-mutant KRAS epitopes in the context of tumors expressing HLA-A2, the most common human HLA-A allele, have remained elusive despite evidence an epitope encompassing such mutations can bind HLA-A2 and induce T cell responses. We report post-translational modifications (PTMs) on this epitope may allow tumor cells to evade immunologic pressure from TCR-T cells. A lysine side chain-methylated KRAS_G12V_ peptide, rather than the unmodified epitope, may be presented in HLA-A2 by tumor cells and impact TCR recognition. Using a novel computationally guided approach, we developed by mutagenesis TCRs that recognize this methylated peptide, enhancing tumor recognition and destruction. Additionally, we identified TCRs with similar functional activity in normal repertoires from primary T cells by stimulation with modified peptide, clonal expansion, and selection. Mechanistically, a gene knockout screen to identify mechanism(s) by which tumor cells methylate/demethylate this epitope unveiled SPT6 as a demethylating protein that could be targeted to improve effectiveness of these new TCRs. Our findings highlight the role of PTMs in immune evasion and suggest identifying and targeting such modifications should make effective ACTs available for a substantially greater range of tumors than the current therapeutic landscape.

**One-sentence summary:** Tumor cell methylation of KRAS_G12V_ epitope in HLA-A2 permits immune evasion, and new TCRs were generated to overcome this with engineered cell therapy.

## Introduction

T cell receptors (TCRs) specific for a tumor-derived MHC class I epitope can be transduced into patient T cells for adoptive cell therapy (ACT) to achieve promising antitumor effects (*1, 2*), and a subset of such TCRs have been advanced into clinical trials (*3–5*). The most desirable TCR-T cell therapies target antigens are tumor-specific, contribute to and/or are essential to the malignant phenotype, and well-presented by the tumor being targeted. Mutant KRAS is the most common driver oncogene in human malignancies (*6*), frequently mutated in pancreatic, colon, and lung cancers. 83% of gain-of-function KRAS mutations result from a change in the glycine amino acid at the 12^th^ residue (*7*). In particular, mutation of glycine to valine, among the most common KRAS mutations in tumors (e.g., in 28.3% of all pancreatic cancers (*8*)), has been postulated to create an epitope (KRAS_G12V_, residues 5-14) presented on tumor cells by HLA-A2 (*9*), the most common Class I allele worldwide (*10*). As tumors often become oncogene-addicted to G12-mutant KRAS (*11*), these mutations are unlikely to be lost or revert to wild type. In pursuit of G12 mutant KRAS as an oncologic target, drugs have recently been developed targeting KRAS_G12C_ by creating a covalent disulfide bond with the mutant cysteine residue. These drugs have yielded promising response rates of 40-50%, but tumors rapidly develop resistance via mechanisms such as structural KRAS mutations outside of G12 that cause steric hindrance of drug activity, amplification of *KRAS* expression, and/or enhancement of downstream oncogenic signaling (*12, 13*); as a result, median progression-free survival remains at about 6.5 months, with no evidence of cures (*14, 15*). Thus, additional strategies for targeting mutant KRAS are needed. The ability to target epitopes containing G12 mutations of KRAS with TCR-T cell therapy would provide an orthogonal strategy for treating mutant KRAS-expressing tumors.

TCRs targeting KRAS epitopes containing a mutation at G12 presented by several HLA alleles including HLA-A3 (*16*), HLA-A11 (*4, 17*), and HLA-C8 (*18*) have been reported to exhibit therapeutic efficacy (*4, 19*). However, accessibility to this therapeutic strategy is limited by the low frequency of these alleles in the US (e.g., 20–24% for HLA-A3 and less frequent for HLA-A11 and HLA-C8) (*20*). In the US, HLA-A2 is the most common allele (47.6% (*21*)); the annual incidence of HLA-A2^+^, mutant KRAS cancers can be estimated to be 93,000 in the United States alone. Thus, targeting an A2-restricted KRAS epitope would enable therapy for a far more substantial fraction of the population. However, defining an A2-restricted KRAS epitope has proven problematic, including difficulties confirming within the sensitivity of mass spectrometry (MS) that an A2-restricted epitope spanning the G12 position is present in the immunopeptidome (*22, 23*). However, KRAS_G12V_ does have epitopes predicted to bind well to HLA-A2, particularly residues 5-14 (KLVVVGAVGV) (*9, 24*). As MS has been able to isolate G12V- containing epitopes and other mutant KRAS epitopes from other HLA alleles (*23*), it has been presumed the HLA-A2 epitope may not be naturally processed and/or efficiently presented by HLA-A2 (*22, 23*).

Several mechanisms may cause defective processing or presentation of a predicted epitope, leading to immune evasion. Examples include differential expression of proteasome subunits that can alter processing of a protein into an epitope (*25*), defective transportation of a processed epitope to the endoplasmic reticulum for binding to MHC I (*26*), and deficiencies in MHC I expression or structure (*27*). Additionally, proteins can undergo post-translational modifications (PTMs), and presentation of an epitope containing a PTM can result in complete (*28*) or partial abrogation of epitope binding or recognition by a TCR reactive to the unmodified epitope. Such changes include oxidation (*28*), nitrosylation (*28*), methylation (*29*), phosphorylation (*30*), and glycosylation (*31*). Some PTMs may also be more prevalent in tumors as a consequence of malignant transformation and enzyme dysregulation (*32, 33*). PTMs do occur in mutated KRAS (*34*), and could interfere with detection of an HLA-A2-presented KRAS epitope by MS due to an altered migration pattern, contributing to the lack of consensus regarding an A2 epitope.

Identifying a targetable epitope with a PTM for KRAS_G12V_ is challenging due to experimental and analytical limitations of MS in detecting PTMs (*35–37*), such as introduction of confounding chemical artifacts (*37*), and PTMs have generally been overlooked when predicting epitopes for therapeutic TCR development (*38*). For the KRAS protein, PTMs are known to occur in both the C-terminal region, which are well-characterized for their signaling functions (*33, 39*), and the N-terminal region with methylation of the lysine-5 side chain (*34*), which is included in the epitopes predicted to be presented by HLA-A2.

In this investigation, we demonstrate that methylation of the lysine-5 side chain in the KRAS_G12V_ epitope that binds to HLA-A2 may interfere with targeting of KRAS_G12V_ tumor cells by TCRs specific for the unmodified epitope. Combining machine learning-based computational approaches to model TCR:pMHC interfaces with high-throughput experimental screens, we developed translationally promising TCRs that can target both methylated and unmodified epitopes, thereby providing broader anti-tumor activity against HLA-A2^+^, KRAS_G12V_^+^ tumor cells. Finally, we uncovered the potential role of a protein canonically known to diminish histone methylation, SPT6 (*40*), in reducing methylation of the KRAS_G12V_ epitope. This may be targetable in tumor cells as a method to enhance immunologic recognition by methylation-targeting TCR-T cells and exemplifies a potential synergistic strategy to inhibit tumor evasion of T cell immunity.

## Results

### Post-translational modifications of the HLA-A2-presented KRAS_G12V_ epitope alter its recognition by MHC class I-restricted TCRs and prevent tumor cell killing

Computational prediction, by NetMHCPan 4.1 (*41*), of HLA-A2-presented KRAS_G12V_ identified several epitopes that specifically contain the G12V mutation (**Fig. S1a**). Among the top ten predicted epitopes encompassing the full mutated KRAS protein, three contained the G12V mutation, and the epitope consisting of residues 5-14 (i.e., KLVVVGAVGV) was predicted to bind with the highest avidity.

For our initial effort to develop HLA-A2-restricted TCRs targeting this peptide, we isolated primary CD8^+^ T cells from peripheral blood mononuclear cells (PBMCs) from HLA-A2^+^ healthy human donors and expanded epitope-reactive clonotypes by stimulation with autologous antigen-presenting cells (APCs) pulsed with a synthetic KRAS_G12V_ 5-14 peptide that lacked any PTMs (**Materials and Methods**). We sorted cells with potentially high affinity for the KRAS_G12V_ peptide as measured by binding limiting concentrations of HLA-A2:KRAS_G12V_(_5-14_) tetramers. From the sorted cells, we then quantitated TCR clonotypes using the ImmunoSEQ assay (Adaptive Biotech) and identified paired TCRa/TCRb sequences using single cell RNA-seq. TCR clonotypes preferentially enriched within the sorted populations were synthesized as codon-optimized P2A-linked constructs of TCRa and TCRb in a lentiviral backbone and used to transduce CD8^+^ T cells. The cells were stimulated with titrated concentrations of KRAS_G12V_(_5-14_) peptide, and the resulting IFNγ expression measured to assess functional avidities for this peptide (**Fig. 1a; Fig. S1b**). Two particularly high affinity TCRs were identified, TCR_19_ and TCR_2_, with the former exhibiting the highest functional avidity. However, in a coculture with the HLA-A2^+^, KRAS_G12V_^+^ pancreatic adenocarcinoma cell line CFPAC1, CD8^+^ T cells transduced with TCR_19_ unexpectedly demonstrated poorer inhibition of tumor growth compared to T cells transduced with TCR_2_ (**Fig. 1b**), whereas TCR_19_ conversely demonstrated better inhibition than TCR_2_ of DAN-G tumor cell growth, which are also HLA-A2^+^ and KRAS_G12V_^+^.

**Fig. 1:**
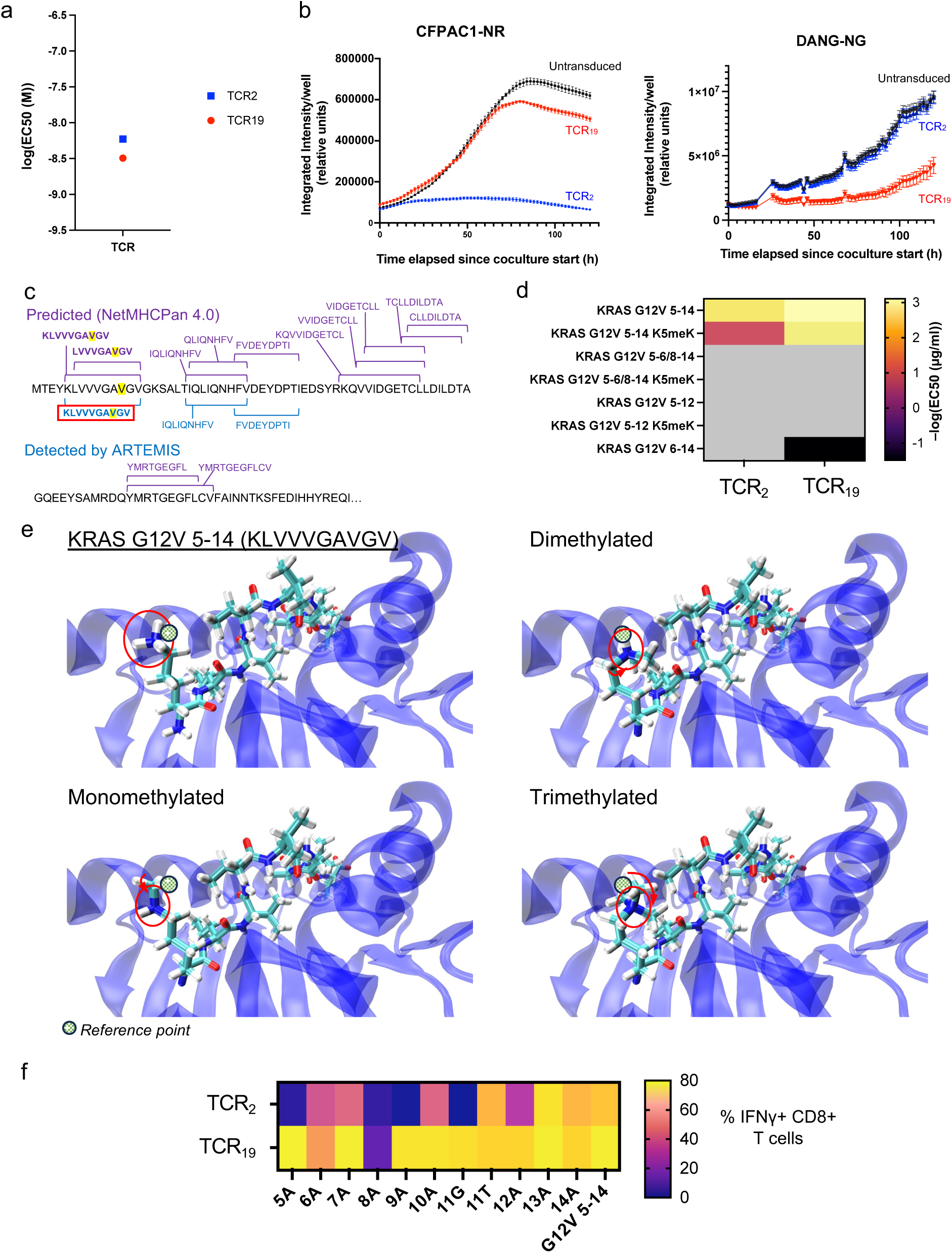
Posttranslational modification on the HLA-A2-presented KRAS_G12V_ epitope alters recognition of the antigen by MHC class I-restricted TCRs and subsequent tumor cell killing. (a) Log_10_(EC_50_ (M)) comparison of recognition of different concentrations of a synthetic KRAS_G12V_ 5-14 peptide (KLVVVGAVGV) presented by HLA-A2 by different MHC class I-restricted TCRs specific for the epitope. Lower values indicate higher functional avidity. (b) Killing of live CFPAC1 and DAN-G HLA- A2^+^ KRAS_G12V_^+^ pancreatic adenocarcinoma cells cocultured with CD8^+^ T cells expressing the TCRs shown in (a) using a 4:1 T cell:tumor cell ratio. Cocultures were recorded by Incucyte imaging. Tumor cells expressed nuclear fluorescent protein; NR: red fluorescent protein; NG: green fluorescent protein. (c) ARTEMIS MS data of peptides eluted from HLA-A2. Data is from the 293F cell line, as commonly used for proteomics assays requiring high protein expression. All cell lines were transduced with an HLA- A2 single chain secreted dimer as well as a KRAS_G12V_ constitutive expression construct to increase presentation and the likelihood of detecting a presented peptide by MS, of KRAS_G12V_-related epitopes. The first 100 amino acids of KRAS_G12V_ are shown, with the G12V mutation highlighted in yellow. Epitopes predicted by NetMHCPan 4.1 are shown in purple; epitopes detected from ARTEMIS are shown in blue. ARTEMIS detection of KRAS_G12V_ 5-14 is outlined in red. (d) Heatmap of –log_10_(EC_50_ (μg/ml)) of CD8^+^ T cells expressing TCR_2_ or TCR_19_ against candidate KRAS_G12V_ epitopes, with and without methylation of the lysine-5 side chain of the epitope, presented by HLA-A2. Higher values indicate higher functional avidity. EC_50_ values were calculated from T cell exposure to peptide concentrations ranging from 1 µg/ml to 10^-5^ µg/ml. Gray squares indicate EC_50_ calculations that lacked a stable fit, e.g., due to lack of response to peptide even at high doses. (e) Rosetta structural modeling of HLA-A2 presenting KRAS_G12V_ 5-14 and its methylated variants. Red circles indicate the amine group of the lysine-5 side chain; arrows indicate its movement with different methylation states. A reference point is drawn at the same position in each image to help illustrate positional changes. (f) Alanine scan of the KRAS_G12V_ epitope and resulting response by CD8^+^ T cells expressing TCR_2_ and TCR_19_. The epitope with each individual residue substituted with alanine (or, for alanine-11, substituted with either glycine or threonine) was presented by antigen-presenting cells to stimulate primary human CD8^+^ T cells expressing a TCR, and subsequent IFNγ expression was measured. Higher IFNγ induction indicates greater tolerance for amino acid substitution at that position. X-axis numbers indicate the residue number on the KRAS protein, and letters indicate the amino acid substitution. A: alanine; G: glycine; T: threonine. Last column is unmodified KRAS_G12V_ 5-14.

The reasons for this discrepancy between TCR functional avidities for KRAS_G12V_(_5-14_) and killing of different tumor cells were unclear. One explanation could be alloreactivity rather than tumor antigen specificity. For killing of CFPAC1 cells, alloreactivity to this cell line by TCR_2_ would not necessarily have been predicted by measuring the dose response of cells with this TCR to peptides presented by another antigen-presenting cell. However, TCR_2_ did not induce a reaction against any of the Class I HLA alleles found on CFPAC1 cells on other targets not expressing mutant KRAS (HLA-A2, A3, B35, B73, C4, C15; **Fig. S1c**). Alternatively, TCR_19_ killing of DAN-G cells could reflect allorecognition, but similar studies did not detect recognition of the alleles on DAN-G by TCR_19_. Thus, alloreactivity was an unlikely explanation. We therefore hypothesized that, while both of these HLA-A2^+^ KRAS_G12V_^+^ tumor cells may be presenting the epitope, one may be presenting it in a modified form, and that there was differential recognition of the modified and unmodified peptide by each TCR.

Before more closely examining the presence of PTMs, we wanted to confirm that the predicted primary sequence, KRAS_G12V_ 5-14, is in fact efficiently generated by proteasomal cleavage and presented by HLA-A2. To accomplish this, we used the ARTEMIS technology, which entails MS-based identification of MHC I epitopes bound to a secreted single-chain dimer (SCD) of the selected HLA I allele (*42*) (**Materials and Methods**). This provides an abundant source of easily purified peptide-bound Class I molecules, can detect the majority of peptides that get endogenously processed and loaded on Class I in the ER, and overcomes the challenge of identifying the limited amount of peptide specifically presented on the cell surface by an individual Class I allele using MS (*42*). We established an ARTEMIS system in the 293F cell line, which supports exceptionally abundant protein expression compared to other mammalian cell lines (*43*). 293F cells were lentivirally transduced to express a secreted HLA-A2 SCD as well as a truncated version of the KRAS_G12V_ protein (amino acids 1–100 to render the protein nonfunctional but preserve the N-terminal portion of the protein that contains the G12V mutation), to increase the availability of KRAS_G12V_-originated peptides that contain the G12V mutation for binding to HLA-A2. Three HLA-A2-presented KRAS peptides were identified by ARTEMIS, including the KRAS_G12V_ 5-14 peptide, which was the only HLA-A2-presented peptide containing the mutated G12V residue (**Fig. 1c**). These results confirmed that the 5-14 epitope is a dominant product of KRAS_G12V_ antigen processing and presentation associated with HLA-A2 and is the correct peptide sequence to target with TCRs.

Given the known dysregulation of methylation processes in mutant KRAS-driven cancers (*32, 33, 44, 45*), and the previously detected lysine-5 side chain methylation of Ras family proteins (*34*), we considered the possibility that the KRAS_G12V_ 5-14 epitope presented by tumor cells contains some level of methylation or other post-translational modification. Having confirmed the primary sequence of the epitope, we hypothesized that DAN-G cells were presenting KRAS_G12V_ 5-14 epitope such that some combination of lysine-5-methylated and unmodified epitope was likely being presented. In turn, the presence of the weakly recognized, modified epitope may have limited the recognition of these tumor cells by TCR_2_ and TCR_19_, thereby limiting tumor killing. However, as 293F cells may not have the same protein methylation machinery active in DAN-G, we attempted to design ARTEMIS in DAN-G cells. Unfortunately, due to the low cell density of cultured DAN-G cells and lower level of protein expression compared to 293-based ARTEMIS cells, we did not detect sufficient levels of peptide to identify the methylated epitope. Therefore, to search for differences in methylation activities that can lead to increased methylation within KRAS in DAN-G cells, we also performed immunoprecipitation of whole-protein KRAS from DAN-G and CFPAC1 cells, followed by liquid chromatography-MS. This revealed that in DAN-G cells, but not in CFPAC1 cells, lysine side chain trimethylation was present at lysine-16 (**Fig. S1d**), representing full replacement of all hydrogens of the amine group by methyl groups. No peptides containing lysine-5 of KRAS were able to be recovered or detected in this data set, likely due to the cleavage of the KRAS protein by trypsin immediately after lysine-5 during sample processing for MS (*46*), producing a short 5-mer peptide that would be difficult to detect with our MS technology. Thus, while we could not directly confirm the presence of a lysine-5-methylated peptide, a pattern of increased KRAS lysine methylation was evident in DAN-G cells. Additionally, previous reports clearly demonstrated methylation of KRAS at lysine residues (*33, 34*) including dimethylation at lysine-5. Lysine side chain methylation may exist as mono-, di-, or trimethylation, with the former two states being intermediate degrees of full methylation.

To determine whether the KRAS_G12V_(_5-14_) peptide with lysine-5 methylation binds to HLA-A2 with comparable affinity as the unmodified peptide, we assessed the stability of HLA-A2:KRAS_G12V_ pMHC complexes with or without lysine-5 side chain methylation. T2 cells are deficient in Transporter Associated with Antigen Processing (TAP) protein and express few properly folded Class I molecules on the cell surface (*47*). Pulsing T2 cells with exogenous peptide can stabilize the conformation of HLA-A2 molecules on the cell surface, resulting in enhanced surface expression that correlates with peptide affinity for HLA-A2 (*47*). We pulsed T2 cells with KRAS_G12V_(_5-14_) with or without methylation and measured the resulting surface expression of HLA-A2 (**Materials and Methods; Fig. S1e**). Compared to untreated control cells, HLA-A2 surface expression was higher when T2 cells were pulsed with the KRAS_G12V_(_5-14_) peptide. Moreover, pulsing with the same peptide containing different degrees of lysine-5 methylation further increased A2 expression, indicating that methylation may enhance the stable presentation of the epitope. Although the unmodified peptide was recognized well by both TCRs, there were large differences in the abilities of the TCRs to recognize the methylated epitope (**Fig. 1d**). Therefore, differential lysine 5-methylation may explain the disparities in the killing of different tumor cells observed with these TCRs.

In addition to the 10-mer KRAS_G12V_ 5-14 sequence detected from ARTEMIS, a 9-mer (LVVVGAVGV) is another predicted epitope containing the G12V mutation that bind HLA-A2 (**Fig. S1a**), and a previous study has also reported peptide splicing to produce an HLA-A2-presented epitope containing residues 5-6/8-14 of KRAS_G12V_ (*24*). To rule out recognition of these alternative epitopes by the TCRs and to test recognition of methylated epitope, we synthesized these epitope variants as well as KRAS_G12V_(_5-14_), including this peptide post-translationally modified at lysine-5. To determine the functional avidities of the TCRs for these variant epitopes, we stimulated T cells expressing TCR_2_ or TCR_19_ with these epitopes as presented by HLA-A2 and measured resulting IFNγ expression (**Fig. 1d**). Indeed, while both TCR_2_ and TCR_19_ recognized unmethylated KRAS_G12V_ 5-14 to similar degrees, only TCR_19_ produced a response to the peptide with methylation of the amine group on the side chain of lysine-5. Other forms of the G12V-containing epitope which had lower predicted binding ability from NetMHCPan 4.1 (residues 6-14 or 5-12), and an alternatively spliced peptide KRAS_G12V_ 5-6/8-14 previously suggested to be processed (*24*), also did not elicit a response from the TCRs. Thus, the DAN- G cell line may be modifying a fraction of KRAS_G12V_ protein or its resulting epitopes with lysine-5 side chain methylation, such that the epitope presented by this tumor cell could not be recognized by TCR_2_. The presence of lysine-5 methylation is consistent with prior findings examining methylation of RAS family proteins, including KRAS (*34*), and is consistent with the known dysregulation of protein methylation in mutant KRAS-driven cancers (*32, 33*).

Based on the evidence supporting contribution of a lysine-5 methylated peptide to TCR recognition of the HLA-A2-presented KRAS_G12V_(_5-14_) epitope, we sought to identify structural changes that may occur as a result of such methylation. Using the Rosetta (*48*) protein structure modeling tool, we introduced monomethylation, dimethylation, and trimethylation to the lysine-5 side chain and evaluated the resulting changes in peptide positioning in the MHC cleft. This computational pipeline indicated that methylation of the lysine-5 side chain within HLA-A2-presented KRAS_G12V_(_5-14_) induces shifts of the positioning of the amine group of this side chain that could affect TCR recognition of the epitope (**Fig. 1e**): in the unmodified peptide, the amine group of the lysine-5 side chain protrudes out of the MHC I cleft potentially into close proximity to the TCR chain, suggesting it may serve as a contact point between the TCR and pMHC complex. Monomethylation leads to displacement of the amine group deeper into the MHC I cleft by the methyl group, which is further pronounced by dimethylation or trimethylation, potentially explaining the observed impact of methylation on the recognition of this pMHC complex by TCR_2_ or TCR_19_.

To evaluate the relative contribution of the lysine-5 residue to binding of each TCR to the HLA- A2-presented KRAS_G12V_ epitope, we performed alanine scans, in which each residue of the HLA-A2- presented epitope is sequentially substituted with the small, neutral alanine amino acid. Each alanine mono-substituted epitope was then used to pulse antigen-presenting cells to stimulate CD8^+^ T cells expressing TCR_2_ or TCR_19_, and IFNγ was then measured to determine if recognition of the whole epitope is impacted by the alanine substitution (*49*), which would imply that the substituted amino acid is a relevant contact point for the TCR. Residue 11 of the 5-14 epitope, which is already alanine, was alternatively substituted with threonine or glycine. At peptide concentrations of 0.1 µg/ml, the analysis revealed that substitution of lysine-5 to alanine reduced the response to the epitope from 67.9% responsive cells to 2.45% for TCR_2_ (**Fig. 1f**), suggesting that TCR_2_ is sensitive to changes at that position of the epitope. The pMHC complex with KRAS_G12V_ epitope, as modeled by Rosetta, also indicates protrusion of the lysine-5 side chain out of the groove of HLA-A2 in the direction where it could interact with a TCR (**Fig. S1f**, which uses the same models as **Fig. 1e** but is rotated for display from a perspective showing this protrusion more clearly). By contrast, the alanine scan for TCR_19_ revealed high tolerance for amino acid substitutions at most positions of the KRAS_G12V_ epitope, which may explain its recognition of methylated lysine-5 but suggesting it would be promiscuous and less suitable as a potentially therapeutic TCR to modify for pursuing improvements (**Fig. 1f**).

### A rationalized mutagenesis approach can be used to generate TCRs that recognize methylated KRAS_G12V_ epitope to improve targeting of tumor cells

Identifying TCRs that can recognize lysine-5 side chain mono-, di-, and/or trimethylation of the KRAS_G12V_ 5-14 peptide alone or in addition to the unmodified version of that peptide could potentially increase the ability to therapeutically target HLA-A2^+^ tumors with the KRAS_G12V_ mutation. To generate new TCRs that can target the peptide with the three different degrees of lysine-5 methylation, we first explored a strategy to mutagenize the CDR3α region of an existing TCR, TCR_2_, which recognizes the unmodified peptide with high functional avidity and specificity for the KRAS_G12V_ epitope, to determine if we could broaden peptide recognition to include methylated variants without acquisition of more general promiscuity.

To determine the region of this TCR most likely to interact with the methylation site, we developed a new computational structural prediction workflow (**Fig. S1g**). We first used Alphafold (*50*), which utilizes deep learning-based approaches to predict the structure of the TCR:pMHC complex without any PTMs, as the latter are not modeled well by Alphafold. We fed the output into Rosetta, which utilizes physics-based energy functions to allow for modeling additional molecular changes to that structure, such as introducing PTMs to the epitope, and then readjusting the structural prediction. Our workflow, performed on the existing TCR_2_ that recognized the unmodified KRAS_G12V_ 5-14 peptide presented by HLA-A2, revealed a core sequence within the CDR3α region most likely to be in close proximity to the lysine-5 side chain and any potential methylation modifications to this residue (**Fig. 2a**).

**Fig. 2:**
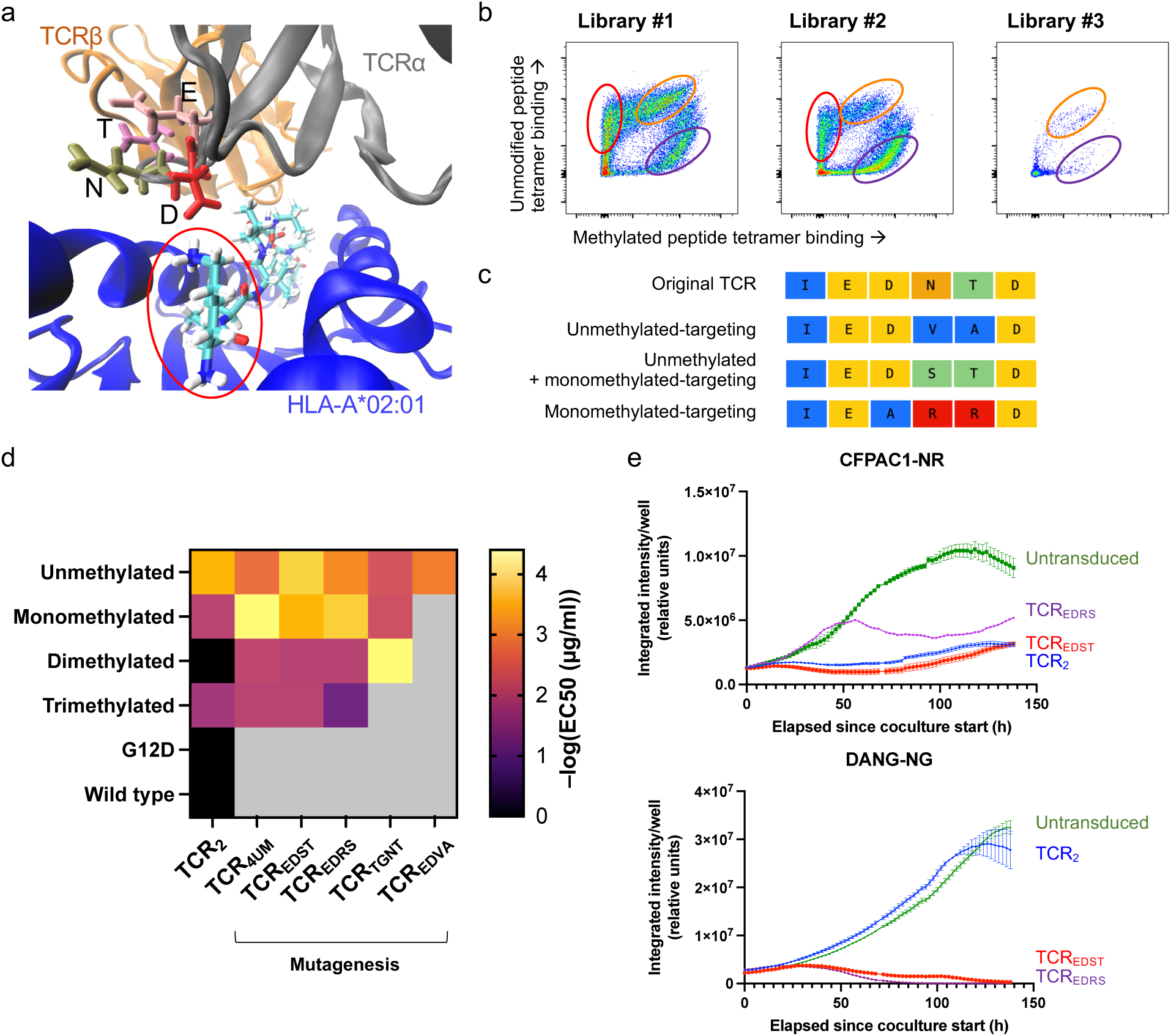
A computationally guided, rationalized mutagenesis library to develop MHC class I-restricted TCRs that recognize methylated epitopes of HLA-A2-presented KRAS_G12V_ and achieve enhanced targeting of HLA-A2^+^ KRAS_G12V_^+^ tumor cell lines. (a) Structural model, as predicted by a combined Alphafold-Rosetta pipeline, of a ‘core’ CDR3α sequence (sequence EDNT) of an existing TCR recognizing KRAS_G12V_ 5-14 and its relation to the lysine-5 side chain (circled in red) of the epitope as presented by HLA-A2. (b) Generation of TCR mutagenesis libraries by mutagenesis of the core CDR3α sequence followed by transduction of the TCRs into Nur77-GFP reporter Jurkat cells and subsequent clonal expansions. Tetramer binding patterns of HLA-A2-KRAS_G12V_ 5-14 peptide with unmodified or methylated lysine-5 side chain at the end of cell stimulation are shown. Each circle indicates a population with a specific pattern of unmethylated vs. methylated peptide tetramer binding that was sorted and sequenced. (c) Examples of mutated CDR3α ‘core’ sequences resulting from sorted clones of the mutagenesis library that exhibit the noted different patterns of PTM peptide recognition. (d) Heatmap of – log_10_(EC_50_ (μg/ml)) of TCRs derived from the mutagenesis library and the original TCR_2_ responding to different degrees of lysine-5 methylation of KRAS_G12V_ 5-14 presented by HLA-A2, as calculated from dose response assays. Gray squares indicate EC_50_ values that lacked a stable fit, e.g., due to limited response to peptide even at high doses. TCR_4UM_ is the 4^th^ TCR clonotype identified from an initial sequencing dataset from sorted Jurkat cells binding to both unmethylated (“U”) and methylated (“M”) tetramers; other TCRs are labeled by changes in the ‘core’ CDR3α sequence (e.g., EDST). (e) Killing of HLA-A2^+^ KRAS_G12V_^+^ CFPAC1 and DAN-G pancreatic adenocarcinoma cells by CD8^+^ T cells expressing selected mutagenized TCRs. Cocultures of T cells and tumor cells were recorded by Incucyte imaging. Tumor cells expressed nuclear fluorescent protein. NR: red fluorescent protein; NG: green fluorescent protein.

Using the structural model as the guide to the most relevant region, we created a rationalized mutagenesis library of this computationally identified core CDR3α sequence. Library-transduced CD8^+^ Jurkat cells were stimulated to detect TCR clonotypes that recognized differently modified variants of the KRAS_G12V_ 5-14 peptide (**Materials and Methods**). Library-transduced T cells were analyzed for binding to unmethylated or lysine-5-methylated KRAS_G12V_(_5-14_) peptide. A broad spectrum of reactivity towards one or both of these peptides was observed, confirming that the computationally determined CDR3α sequence plays a key role in recognition of the putative lysine-5 methylation site by TCR_2_. We identified several distinct clonotypes able to bind to only the unmethylated peptide (e.g., TCR_EDVA_), to a combination of unmethylated and mono/di/trimethylated epitopes (e.g., TCR_4UM_, TCR_EDST_, TCR_EDRS_), or to only methylated peptides (TCR_TGNT_; **Fig. 2b–c**), which were sequenced to identify the corresponding mutant variants. New constructs were then generated expressing these mutant variant TCRs. A dose response assay with CD8^+^ T cells transduced with each mutant variant was performed using the unmethylated and methylated KRAS_G12V_ epitopes individually. Some mutagenized TCRs, when compared to TCR_2_, had improved recognition of the methylated epitope (**Fig. 2d**). A dose response assay revealed no significant acquisition of recognition of wild type KRAS 5-14 or the alternative KRAS_G12D_ 5- 14 epitope by the mutagenized TCRs compared to TCR_2_ (**Fig. S2a**). Thus, changes in amino acids within only this computationally identified core CDR3α sequence were able to alter the TCR’s pattern of recognition for peptides with PTMs (**Fig. S3a–b**) without altering recognition of non-G12V KRAS epitopes. For TCR_EDST_, labeled as such for the mutagenized ‘core’ CDR3α sequence, the amino acids in closest proximity to the lysine-5 side chain of the HLA-A2-presented KRAS_G12V_ epitope were aspartate and serine. These amino acids are negatively charged and polar, respectively, allowing for electrostatic or hydrogen bonding interactions with the lysine-5 side chain amine group of the KRAS_G12V_ epitope. While methylation introduces a hydrophobic methyl group, the polar nature of the amino acids on this TCR may still allow for binding to the lysine side chain.

We then tested whether CD8^+^ T cells expressing the TCR variants with enhanced binding to methylated peptides could efficiently recognize and kill KRAS_G12V_-expressing tumor cells. In coculture with KRAS_G12V_^+^ HLA-A2^+^ tumor cells, CD8^+^ T cells expressing TCR_EDST_ or TCR_EDRS_, which recognize unmethylated and methylated KRAS_G12V_ epitopes, demonstrated improved killing of DAN-G cells, which likely presents methylated and unmethylated KRAS_G12V_(_5-14_), and retained similar killing of CFPAC-1 cells compared to TCR_2_ (**Fig. 2e**). Consistent with the lack of recognition of wild type KRAS_5-14_ or KRAS_G12D_(_5-14_) in the peptide dose response assay, these TCRs did not inhibit growth of HeLa cells, which express only wild type KRAS_5-14_, or of Panc1 cells, which express KRAS_G12D_ (**Fig. S2b**). Taken together, mutagenesis of the originally naturally isolated TCR_2_ yielded new TCRs recognizing different combinations of unmodified and varyingly methylated KRAS_G12V_ epitopes without increasing “off-target” recognition of G12D or wild type epitopes of KRAS.

A critical concern with TCRs deployed for ACT is that specificity needs to be limited only to the intended targets to avoid off-target toxicities, which is of increased concern after a naturally isolated TCR is mutagenized. To further evaluate the specificity of the selected TCRs, we again performed alanine scans of the target epitope for recognition by the mutagenized TCRs in Jurkat reporter cells expressing CD8ab, TCR_2_ or mutagenized TCR (TCR_4UM_, TCR_EDST_, TCR_EDRS_, TCR_TGNT_, or TCR_EDVA_), and the Nur77- GFP reporter gene as a specific marker of TCR signaling(*51*). After pulsing with 0.1 µg/ml peptide concentrations (∼106 nM for this epitope), a concentration high enough to induce complete responses to the original KRAS_G12V_ (_5-14_) epitope, and with tolerance to a change defined as retaining Nur77-GFP reporter expression in >35% of cells that respond to the original epitope, there were distinct positions common to both the unmethylated and methylated versions of the epitope that tolerated amino acid replacements and remained stimulatory (**Fig. S2c**). For TCR_2_, TCR_EDST_, and TCR_EDST_, the overall tolerance pattern was K-x-x-V-V-x-A-x-x-x, with x signifying tolerance for an alanine substitution at that position; among these three TCRs, TCR_EDST_ exhibited the lowest tolerances. Compared to these TCRs, TCR_4UM_ demonstrated greater tolerance for substitutions, and TCR_TGNT_ and TCR_EDVA_ showed lower tolerances but demonstrated less effective tumor control. Thus, TCR_4UM_, TCR_TGNT_, and TCR_EDVA_ were determined to be less suitable as candidate therapeutic TCRs. Notably, while some TCRs could recognize both unmethylated and methylated versions of the lysine-5 side chain, replacement of lysine-5 with alanine led to a loss of recognition of the presented peptide. The diminution of recognition was greater than observed with substitutions at other peptide positions, suggesting this lysine amino acid remained a major contact point for the mutagenized TCRs as it did for TCR_2_. A scan of the human peptidome for epitopes that accommodated any substitution at these permissive residues yielded PLD3_396-405_, MIGA1_74- 83_, NAGPA_276-285_, K/N/HRAS_5-14_, RSLBB_35-44_, CFA61_23-32/607-616/667-676_, CUL9_635-644/745-754_, CLCN6_80-89_, and MACOI_33-42_ as potential off-target hits. These peptides were synthesized and used to stimulate T cells expressing the TCR. Relative to HLA-A2-presented KRAS_G12V_ 5-14, all of the selected TCRs exhibited reduced responses against these peptides at 0.1 µg/ml concentration (**Fig. S2d**), suggesting the mutagenized TCRs had not acquired significant promiscuity compared to the original TCR_2_. Additionally, for TCR_2_ and TCR_EDST_, at more physiologic concentrations of 0.01 µg/ml (∼10.6 nM), these peptides elicited response rates of <6% compared to 18–19% for the KRAS_G12V_ epitope, as measured by percentage of cells expressing IFNγ (**Fig. S2e**).

### Stimulation of primary T cells with selected peptides followed by clonal expansion reveals TCRs from healthy donor repertoires that recognize methylated KRAS_G12V_ epitope and can target tumor cells

The TCRs generated from the mutagenesis approach were able to specifically target and effectively kill HLA-A2 tumor cell lines expressing KRAS_G12V_ with PTMs. However, such TCRs have not been naturally vetted by the thymic selection process, and thus may pose a risk for cross-reactivity with other peptides from the human proteome despite the safety testing based on the alanine scan. Thus, to identify TCRs from normal peripheral repertoires potentially capable of binding to the methylated KRAS epitopes, primary human T cells from healthy donor PBMCs were stimulated with KRAS_G12V_ peptides including unmethylated, mono-, di-, and tri-methylated lysine-5, and responding T cells were sort purified based on specific binding to individual HLA-A2:PTM-containing KRAS_G12V_ 5-14 tetramers at limiting peptide concentrations. Quantitative TCR repertoire analysis was performed on the purified populations, and highly enriched TCR clonotypes were synthesized and expressed in primary CD8^+^ T cells (**Materials and Methods; Fig. S4a, b**).

As measured by IFNγ expression by TCR-transduced T cells after exposure to unmethylated and methylated KRAS_G12V_ epitopes presented by HLA-A2, these TCRs exhibited dose response patterns specific to the G12V mutation, responded to methylated and/or unmethylated peptides, and did not recognize wild type or G12D mutant epitopes (**Fig. 3a**). The functional avidities of these TCRs, TCR_A2UoM1-1_, TCR_A2UoM1-2_, and TCR_A2UoM1-4_, for the unmethylated and monomethylated peptides were comparable to those of the mutagenesis-derived TCRs, in particular TCR_EDST_ which exhibited superior killing of DAN-G cells and strong recognition of unmethylated and methylated KRAS_G12V_ epitopes. Among the TCRs that exhibited the strongest recognition of methylated and unmodified peptides, coculture with tumor cells of CD8^+^ T cells expressing TCR_A2UoM1-1_ demonstrated superior killing compared to T cells expressing TCR_2_ of the HLA-A2^+^ KRAS_G12V_^+^ DAN-G tumor cell line, inferred to present methylated and unmethylated KRAS_G12V_ 5-14 (**Fig. 3b**). TCR_A2UoM1-1_ also exhibited similar killing as the other TCRs against CFPAC1, which presumably presents unmethylated KRAS_G12V_ 5-14. TCR_A2UoM1-1_ did not recognize titrated doses of HLA-A2-presented wild type KRAS 5-14 or KRAS_G12D_ 5- 14 (**Fig. S4c**) but did exhibit some inhibition of HLA-A2^+^ tumor cell lines expressing either wild type or G12D KRAS (**Fig. S4d**), suggesting that there may be some broader targeting. However, an alanine scan with TCR_A2UoM1-1_ demonstrated very limited tolerance for amino acid substitutions within the KRAS_G12V_ 5-14 epitope (**Fig. S4e**), such that the only potential match in the human peptidome (using the same criteria as our previous alanine scans) was EPIPL_1948-1957_. A dose response assay performed with this peptide revealed no recognition by this TCR (**Fig. S4f**), affirming its highly selective specificity.

**Fig. 3:**
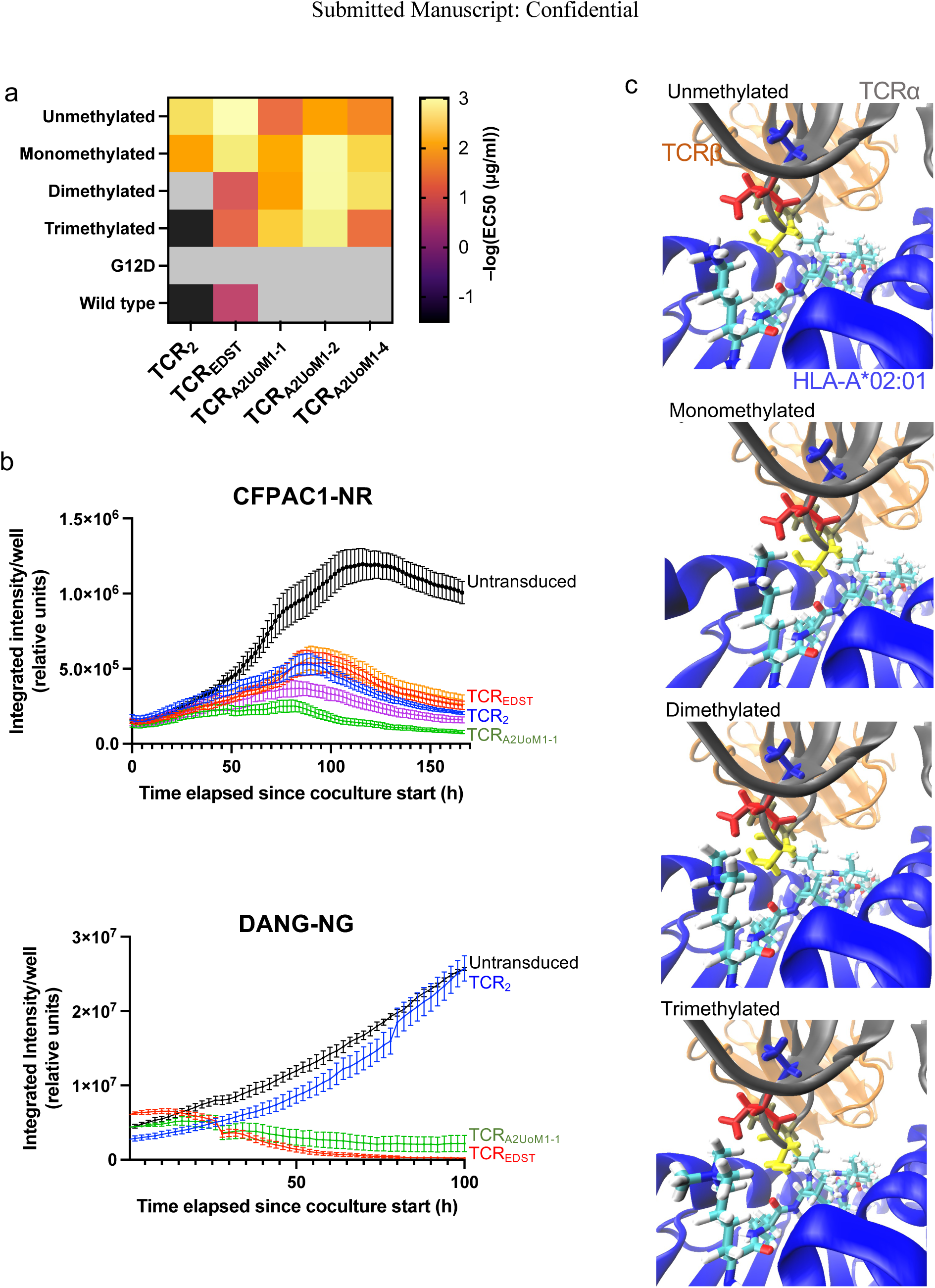
Stimulation of primary human T cells with unmethylated and methylated HLA-A2-presented KRAS_G12V_ to generate MHC class I-restricted TCRs from a normal repertoire that recognize methylated epitopes. (a) Heatmap of –log_10_(EC_50_ (μg/ml)) of selected TCRs responding to the unmethylated and methylated epitopes presented by HLA-A2, as calculated from dose response assays. Higher values indicate higher functional avidity. Gray squares indicate EC_50_ values that lacked a stable fit, e.g., due to limited response to peptide even at high doses. (b) Killing of HLA-A2^+^ KRAS_G12V_^+^ CFPAC1 and DAN-G pancreatic adenocarcinoma cells by CD8^+^ T cells expressing selected TCRs. Cocultures of fluorescently labeled tumor cells and TCR-T cells were recorded by Incucyte imaging. Tumor cells expressed nuclear fluorescent protein; NR: red fluorescent protein; NG: green fluorescent protein. (c) Alphafold-Rosetta visualizations of TCR:pMHC complexes for TCR_A2UoM1-1_, isolated from expansion of responding primary human CD8^+^ T cells from normal repertoires. Structures were generated for unmethylated and mono/di/trimethylated versions of the epitope, presented by HLA-A2. The four CDR3α amino acids in closest proximity to the lysine-5 side chain of KRAS_G12V_ are shown with individual molecular bonds.

Structural modeling of TCR_A2UoM1-1_ revealed a CDR3α sequence that had amino acids aspartate and serine in closest proximity to the lysine-5 side chain including the differently methylated versions (**Fig. 3c**). These TCR amino acids are identical to what was identified in the mutagenized TCR_EDST_, and thus may allow for close interaction with both methylated and unmethylated lysine-5 of the KRAS_G12V_ epitope by both TCRs.

### SPT6 is involved in the mechanism for demethylation of lysine-5 of the KRAS_G12V_ epitope in tumor cells and may be a target for creating presentation of a stable epitope with a methylation-specific TCR

We next sought to determine potential genes involved in KRAS_G12V_ epitope methylation. To address this, we performed a CRISPR-Cas9 library screen targeting genes canonically known to be involved in DNA or protein methylation (**Materials and Methods; Fig. 4a**). The library was transduced into the DAN-G cell line, which is inferred to express the methylated KRAS_G12V_(_5-14_) epitope and is recognized by TCRs that have specificity for the methylated epitope but not effectively by TCRs only specific for the unmodified epitope. Tumor cells were collected before and after co-culture with methylation-responsive TCR-T cells (TCR_EDST_) and TCR-T cells that preferentially target unmodified KRAS (TCR_2_), and the frequencies of each gRNA in surviving tumor cells were determined. TCR_EDST_ was chosen as the methylation-sensitive TCR for this comparison because it differs from TCR_2_ only in its ability to recognize methylated lysine-5. This analysis revealed reduced recovery of certain gene knockouts when cocultured with TCR_EDST_, but not when cocultured with TCR_2_, suggesting that the knocked-out gene conferred a survival disadvantage for tumor cells that might reflect increased presentation of the methylated epitope. In contrast, a higher frequency of gRNAs targeting a given gene in the same comparison of cocultures would mean that the knockout conferred a survival advantage, thereby providing a means of immunological escape from the TCR_EDST_-T cells by diminishing presentation of the methylated epitope.

**Fig. 4:**
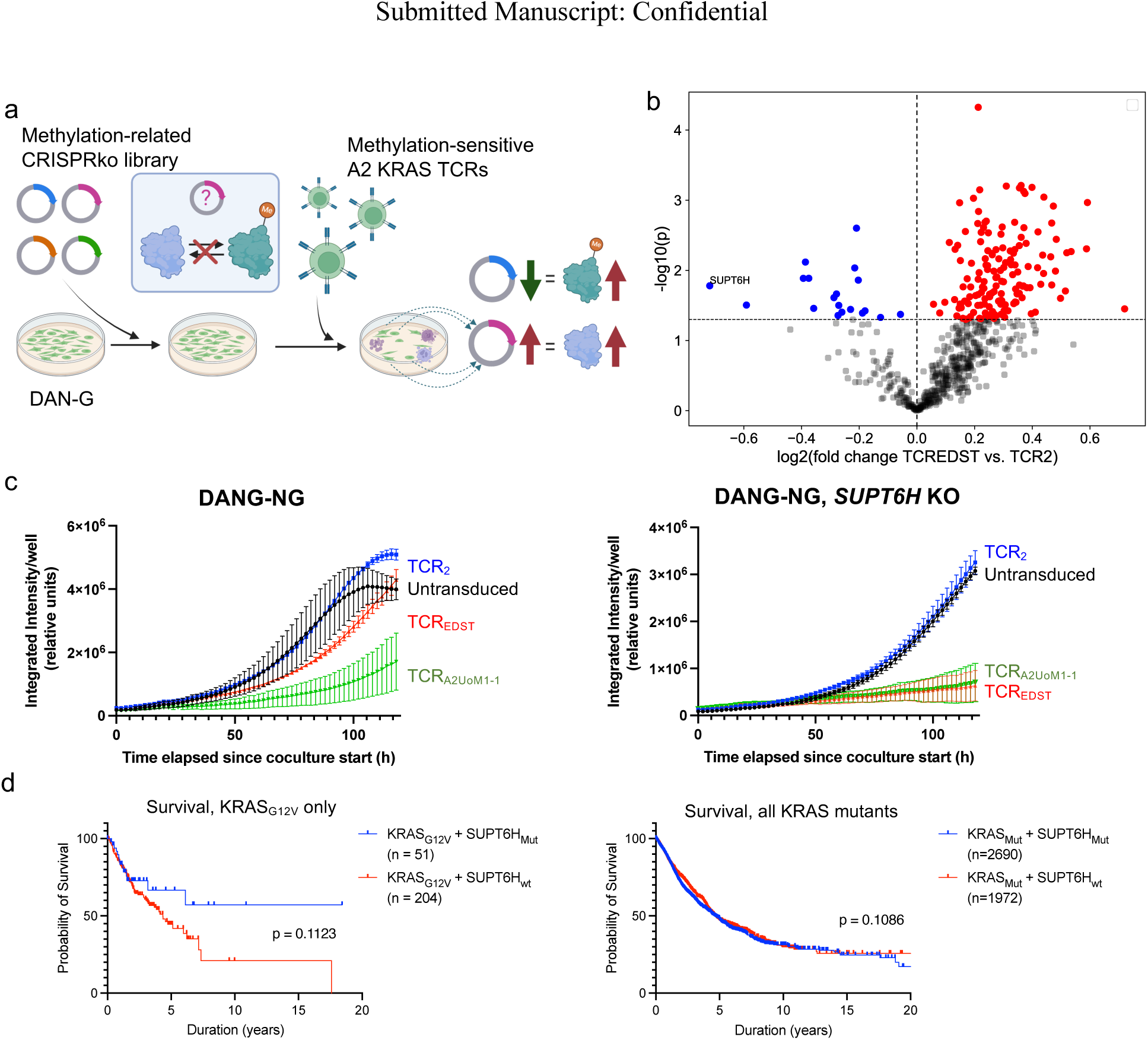
A gene knockout screen for methylation-related genes identifies SPT6 (*SUPT6H*) as a participant in the demethylation of the HLA-A2-presented KRAS_G12V_ epitope. (a) Overall workflow: DAN-G pancreatic adenocarcinoma cells were transduced with a CRISPR knockout library targeting methylation-related genes, and then cocultured with CD8^+^ T cells expressing either TCR_2_ or mutagenized TCR_EDST_, which respectively have weak or strong recognition of methylated KRAS_G12V_ epitopes. Tumor cells surviving at the end of coculture were sequenced to identify gRNA enrichment or depletion associated with immune evasion. (b) Volcano plot summarizing log_2_(fold change) of gRNA read frequencies targeting individual genes recovered after coculture of DAN-G cells with T cells expressing TCR_EDST_ versus TCR_2_. The threshold of significance was set at log_10_(*p*) < 1.2 (*p* < 0.05) and is denoted by the horizontal dashed line. Mean counts of all gRNAs per gene were used to calculate fold change values. The data point for gRNAs targeting *SUPT6H* is specifically labeled. The knockouts depleted or enriched after cocultures using TCR_EDST_ versus TCR_2_ are colored blue and red, respectively. (c) Killing of DAN-G cells with or without *SUPT6H* knockout in coculture with selected TCR-T cells (untransduced or expressing TCR_2_, TCR_EDST_, or TCR_A2UoM1-1_). Cocultures of T cells and tumor cells were recorded by Incucyte imaging. Tumor cells expressed green nuclear fluorescent protein (“NG”). (d) Kaplan-Meier survival plots for individuals who have tumors with KRAS mutations, using public TCGA data. Left: only tumors with the KRAS_G12V_ mutant, with or without concurrent *SUPT6H* mutation. Right: tumors with any KRAS mutant, with or without a concurrent *SUPT6H* mutation.

In the gene knockout screen, DAN-G cells with *SUPT6H* knockout (encoding the SPT6 protein) were the single most depleted knockout when comparing tumor cells cocultured with the methylation-targeting TCR_EDST_-T cell to the non-methylation-targeting TCR_2_-T cell coculture (**Fig. 4b**). This suggests that SPT6 expression decreased methylation of this epitope. SPT6 is canonically known as a negative regulator of histone lysine methylation (*40*). From our data, SPT6 appears to have a non-canonical role as an inhibitor of lysine methylation on the KRAS_G12V_ epitope.

To further investigate the enhanced effectiveness of methylation-targeting TCR-T cells specifically against SPT6-deficient DAN-G cells, we created a *SUPT6H* knockout DAN-G cell line. Successful knockout was verified via long-read sequencing of the targeted genomic region (**Fig. S5a**). Coculture of the *SUPT6H*-knockout DAN-G cells with CD8^+^ T cells transduced either with TCR_EDST_, which targets both unmethylated and methylated KRAS_G12V_ epitope, or with TCR_2_, which only effectively targets the unmethylated KRAS_G12V_ epitope, affirmed that knockout of *SUPT6H* increased susceptibility to killing by T cells expressing either TCR_EDST_ or TCR_A2UoM1-1_ which also targets the methylated epitope (**Fig. 4c**). Additionally, the knockout did not increase killing by T cells expressing the methylation-unresponsive TCR_2_. Relative to the original DAN-G cells, *SUPT6H* knockout did not increase HLA-A2 expression (**Fig. S5b**) and did not change KRAS expression levels (**Fig. S5c**), indicating that the effect of *SUPT6H* knockout was not due to changes in total epitope presentation levels but rather due to changes in the PTM composition of the epitope.

Data from The Cancer Genome Atlas (TCGA) show that loss-of-function mutations of *SUPT6H* specifically within the KRAS_G12V_ mutant cancer patient population lead to increased survival compared to the *SUPT6H*-mutant patient population as a whole (**Fig. 4d**). This mutation-specific behavior may suggest that the methylated KRAS_G12V_ epitope is more spontaneously immunogenic in the context of a naturally occurring TCR repertoire, which would be consistent with the more stabilized cell surface expression of the HLA-A2:KRAS_G12V_ pMHC complex when the epitope is methylated compared to unmethylated, as we described earlier (**Fig. S1e**).

## Discussion

Development of effective ACT using TCRs requires accurate identification of the epitopes specifically presented by the targeted tumor, which can become increasingly difficult in the context of PTMs. In this study, we have demonstrated, using two approaches, the ability to isolate/generate TCRs that recognize and target PTMs that can alter the immunogenicity of a tumor epitope in a therapeutically consequential manner. Deep learning-guided mutagenesis made it feasible to refine a potentially promising TCR targeting an HLA-A2-presented KRAS_G12V_ epitope into one able to target both post-translationally modified and unmodified variants of the epitope, thereby achieving improved killing of HLA-A2^+^, KRAS_G12V_ tumor cell types. We also generated TCRs that can achieve similar activity by selectively expanding rare clonotypes reactive against modified and unmodified KRAS_G12V_ peptide variants from healthy donor-derived primary T cell populations, thereby selecting for TCRs that have undergone the thymic central tolerance process, meaning they are less likely to react against self-antigens and may offer a safer therapeutic profile. For tumor cells that might present the unmodified KRAS_G12V_ epitope together with a mix of variants with different degrees of a lysine-5 methylation, our data suggests that therapeutic TCRs designed or selected to target a wide range of these variants should prove more effective at killing a larger subset of KRAS_G12V_-expressing tumor cells, and preclude immune evasion driven by altered or variable KRAS lysine methylation.

Computational prediction of TCR specificity against pMHC complexes has recently experienced significant improvements (*52*), but purely computational approaches still require experimental validation to determine whether TCR specificity is sufficiently high to be safe and effective for antitumor therapy. Predicting the three-dimensional architecture between a pMHC complex and TCR has the potential to provide insights for experimentally modifying and improving the affinity/specificity of the TCR for the pMHC based on proposed contact points, orientation of a peptide within the pMHC, and orientation of the TCR to that complex. However, the structure of the TCR:pMHC complex has historically been difficult to accurately determine computationally due to the still somewhat limited number of experimentally validated structures of TCR:pMHC interactions from which to base new predictions, the bias of prediction models to overfit based on existing common epitopes, and the lack of negative control data (*53, 54*). Recent innovations in computational protein structure prediction, including Alphafold, have allowed for improved accuracy (*53*). Nonetheless, Alphafold does not yet support the structural prediction of changes resulting from PTMs due to the lack of available structural data for training that includes PTMs (*55*). Alternatively, Rosetta utilizes physics-based energy functions to optimize protein structures, which better accommodates integration of PTMs into protein structure prediction (*56*). Thus, in this investigation, we combined these two pipelines into one prediction to yield a more precise guide that accurately identified pMHC contact sites within the TCR CDR3 sequences that could subsequently be modified through saturation mutagenesis to specifically enhance TCR association with post-translationally modified peptide within the pMHC complex. This strategy allowed us to develop new TCRs that more effectively target HLA-A2^+^ KRAS_G12V_^+^ tumors, while maintaining TCR specificity for mutant KRAS.

Developments in MS technology, including newer technology with greater sensitivity, have allowed for its use to identify MHC class I-restricted T cell epitopes (*35, 42*). We used such MS to characterize the primary sequence of the HLA-A2-presented KRAS_G12V_-containing epitope in this study, analogous to previous studies that identified mutant KRAS epitopes presented by other MHC I alleles (*4, 9, 19*). These MS-based studies have been important for the development of potentially therapeutic TCRs for mutant KRAS tumors. However, MS can remain unreliable for detecting PTMs. The large size of the immunopeptidome presented by the multiple MHC I and II alleles expressed by a tumor, the often-reduced level of MHC expression by tumor cells, the low yield fraction of protein capture, and low expression of some proteins of interest can result in low and inadequate sensitivity for detection of a specific epitope and any associated PTMs (*35*). As a result, MS-based workflows can benefit from maximizing protein expression and MHC capture as we did with ARTEMIS in this study. Additionally, the introduction of chemical artifacts during sample processing, such as oxidation (*37*), may confound whether a detected PTM is genuine or not, and PTMs that are not suspected prior to analysis to be on a peptide may result in missing the detection of that peptide (*36*). To bypass these difficulties associated with epitope detection and probe for a specific PTM on a presented epitope, TCRs with distinct PTM- peptide recognition patterns may be used as an alternative to MS, as these TCRs will elicit T cell activation in response to a target presenting distinct PTMs and can serve as a very sensitive assay for the presence or absence of the PTM.

Based on the results of our efforts to generate TCRs targeting HLA-A2-presented KRAS_G12V_, TCR_EDST_ and TCR_A2UoM1-1_ appear to be two promising candidates for further development as therapeutic TCRs. The former was generated by mutagenesis of an existing TCR that naturally recognized an unmodified epitope derived from mutant KRAS, whereas the latter was identified by selection from a healthy donor’s TCR repertoire. The mutagenesis approach allows for generation of TCRs that may not be present in normal repertoires and that have increased affinity for the target. However, because these TCR sequences are mutated from a TCR naturally existing in the repertoire, they have not been naturally screened during development against the self-immunopeptidome, and pose an elevated risk of off-target activity in a therapeutic setting. Thorough testing for TCR recognition of cross-reactive targets *in vitro*, as we have initiated with alanine scans followed by identification and testing of potential non-KRAS_G12V_ targets in the proteome, should help to mitigate this problem. Additionally, such TCRs may be useful as probes to identify the presence of particular PTMs on epitopes presented by different tumors. TCR discovery from the repertoire of a healthy donor that has already been centrally thymically tolerated is theoretically safer than TCRs generated by mutagenesis, but can be limited if the desired TCR clonotypes are particularly rare within the repertoire or only of low affinity. Overall, we show that either approach is feasible to generate potentially effective, therapeutic TCRs able to target a modified and/or unmodified tumor antigen of interest.

Our findings with the histone methylation inhibitor SPT6 (*SUPT6H*) provide further evidence that PTMs may be introduced by potentially non-canonical processes (*32, 57*), analogous to the non-canonical methylation activity of SMYD3 that has been shown to impact other components of the KRAS signaling pathway (*32*). From our findings, *SUPT6H* knockout in tumor cells enhances the efficacy of KRAS-targeting, PTM-reactive, TCR-T cells. Diminishing SPT6 activity may provide another opportunity to increase tumor immunogenicity, such as by combining these TCR-T cells with a small molecule inhibitor of SPT6.

Our investigation into lysine methylation on KRAS_G12V_ antigen revealed an intriguing role of PTMs in changing recognition of tumor cells by cytotoxic T cells and provided a strategy to mitigate the effects of PTMs to produce more consistently performing therapeutic TCRs. PTMs of tumor-associated antigens have previously been implicated in altering immunogenicity of tumor cells in several tumor types (*58*), and we build upon this by describing a strategy for the development of TCRs to specifically target them. This strategy paves the way for developing TCRs that specifically target PTMs, offering a novel avenue for personalized immunotherapy. Moreover, we have identified TCRs that can target and kill tumor cells with mutated KRAS_G12V_ and that are restricted to the most common human HLA allele, which therefore has significant direct potential for clinical development. The widespread expression of the KRAS_G12V_ mutation across numerous different tumor types, including many solid tumors, means these TCRs may allow for a substantially broadened therapeutic scope of ACT to combat these otherwise difficult-to-treat tumors, including a wide variety of solid tumors.

The roles of PTMs found on tumor antigens outside of altering tumor cell immunogenicity are likely broad. In the case of HLA-A2-presented KRAS_G12V_, lysine-5 methylation in KRAS has been speculated to affect interactions with relevant signaling proteins in the Ras family pathway signaling (*34*). Characterizing the mechanistic reasons for and consequences of PTMs on tumor antigens should grant additional insight into possible molecular pathways that could be targeted synergistically with, or independently of, immunotherapies targeting these antigens.

## Materials and Methods

### Computational modeling of TCR:pMHC complexes

We used Alphafold to generate an initial structural model comprising a given TCR and pMHC complex with unmodified KRAS epitope. Using the resulting model of the TCR:pMHC complex not yet including any PTMs, we used PyRosetta (*48*) 2023.49 to introduce relevant PTMs to the epitope, then reoptimize the binding energy of the TCR:pMHC complex with the PTMs under consideration to create the final structural model. Subsequent visualizations were generated using VMD (*59*). For modeling only the pMHC complex, we used PyRosetta to take a pMHC complex with known structure, consisting of HLA-A2 presenting a peptide with a sequence similar to KRAS_G12V_ 5-14, LLFGYPVAV (PDB: 1QSF (*60*)), mutated the peptide to KRAS_G12V_ 5-14 with or without PTMs, and reoptimized the binding energy of the resulting pMHC complex.

### Generation and use of lentiviral constructs for TCRs

We created lentiviral transfer plasmids expressing our TCRs, using pRRLSIN as a basis. We digested this plasmid with AscI and PstI restriction enzymes (New England Biolabs), with recognition sites being introduced at appropriate locations beforehand via site-directed mutagenesis, and gel-extracted the backbone. We then performed Gibson assembly using the Gibson Assembly Kit (New England Biolabs) on the digested backbone and synthesized DNA blocks (Codex DNA; Integrated DNA Technologies) encoding codon-optimized versions of our selected TCR sequences. The Gibson assembly products were transformed into chemically competent 5-alpha *E. coli* bacteria (New England Biolabs). The final construct thus contained a single open reading frame for the TCR, composed of the TCRβ chain, P2A, and TCRα chain and driven by the MSCV U3 promoter. To verify sequences of resulting clones, we used the full plasmid sequencing service from Primordium Labs.

To generate lentivirus for TCR expression, we cultured 293T cells in RPMI 1640 media (Thermo Fisher) with 5% FBS (Cytiva), 5% human serum (Bloodworks Northwest), 0.6% MEM Non-Essential Amino Acids solution (Thermo Fisher), 4 mM L-glutamine (Gibco), and 100 U/ml penicillin-streptomycin (Gibco). To generate lentivirus from the above constructs, we transfected the 293T cells at 3 million cells/10 cm dish with Effectene transfection reagent (Qiagen) following manufacturer guidelines, using 1.5 μg of an individual TCR construct, 0.5 μg of pMD2.G (Addgene #12259), 1 μg of pMDLg/pRRE (Addgene #12251), and 1 μg of pRSV-Rev (Addgene #12253). 1 d after transfection, we changed media, removing transfection reagent; 1 d following that, we filtered culture supernatant through 0.45 μm PVDF filters. The filtered product was stored at –80°C until used in TCR-T cell generation.

### TCR-T cell generation

We isolated CD8^+^ T cells from primary human donor PBMCs using the EasySep Human CD8^+^ T cell Isolation Kit (STEMCELL Technologies) and activated them overnight using anti-CD3/CD28 DynaBeads (Thermo Fisher). Afterward, we transduced these T cells with lentivirus to express an engineered TCR targeting HLA-A2-presented KRAS_G12V_ epitope, including spinfection at 1,055 *g* for 90 minutes at 30°C. Cells were grown in “CTL media”: RPMI 1640 media (Thermo Fisher) with 10% human serum (Bloodworks Northwest), 4 mM L-glutamine (Gibco), 100 U/ml penicillin-streptomycin (Gibco), and 50 IU/ml IL-2 (Prometheus; obtained via University of Washington Pharmacy, DOH license #DRRS.FX.60550751), with medium changes every 2-3 d. Cells were sorted for peptide-MHC tetramer binding 6 d after transduction. These cells were expanded via rapid expansion protocol for 8–14 d as described previously(*61*). At the end of expansion, the cells were used for functional assays.

### MS of KRAS protein and ARTEMIS

We performed ARTEMIS following previously described procedure(*42*) on DAN-G human pancreatic adenocarcinoma and 293F cell lines. Briefly, we transduced these cell lines with a construct expressing secreted, His-tagged HLA-A2 single chain dimer (SCD) with mCherry and a construct for constitutive expression of the first 100 amino acids of human KRAS_G12V_, based on Lenti-mCherry-G12V- KRAS-IRES-Blast (Addgene #153336). Cells were sorted for SCD expression via mCherry and further selected for the truncated KRAS_G12V_ expression via blasticidin at 10 μg/ml for 7 d. We expanded these cells and harvested culture supernatant for two weeks. We filtered out debris from this supernatant using a 0.6 µm filter, captured HLA-A2 SCD-peptide complex with Ni-NTA agarose beads (Invitrogen) in imidazole equilibration buffer (10mM imidazole, 300 mM NaCl, 50 mM NaH_2_PO_4_, pH 8), and eluted the bound peptides in a denaturing buffer (5M guanidine-HCl, 250 mM NaCl, 50 mM NaH_2_PO_4_, 1 mM DTT, pH 8).

For whole KRAS protein, we performed cell lysis on HLA-A2^+^ tumor cell lines to collect protein as described previously(*62*). From the resulting protein extract, we performed immunoprecipitation for KRAS using MS-compatible magnetic immunoprecipitation kit (Pierce) and anti-KRAS antibody (12063- 1-AP; Thermo Fisher), following manufacturer guidelines. The immunoprecipitated protein was run through SDS-PAGE, and a protein band at ∼21 kDa size was excised and used for liquid chromatography-MS.

### Discovery of new TCRs via mutagenesis of an existing TCR targeting HLA-A2-presented KRAS_G12V_ 5-14

To generate rationalized mutagenesis libraries of TCR_2_ targeting the sequence within CDR3α as selected from the structural prediction, we performed Gibson assembly to combine synthesized saturation mutagenesis DNA blocks replacing the identified ‘core’ residues of CDR3α with the remainder of the TCR_2_ construct. We transduced these TCR libraries into populations of Jurkat cells expressing CD8 and the Nur77-green fluorescent protein (GFP) reporter gene, which specifically permits measurement of TCR signaling(*51*). These cells were grown in CTL media with the addition of KRAS_G12V_ 5-14, with or without methylation of the lysine-5 side chain, and presented by HLA-A2. After two weeks, we sorted these cells for binding to tetramers of HLA-A2 with unmethylated or methylated KRAS_G12V_ epitope and amplified CDR3α sequences from sorted cells. Amplicons were sequenced by next-generation sequencing on an Illumina MiSeq v2 flow cell with paired-end 150 bp reads.

To process sequencing results, we used custom code, trimming adapters and using Bowtie2 to align reads to a reference sequence for the CDR3α region of TCR_2_. We then scanned for variations from the reference sequence and compared the frequencies of individual sequence variants between sorted populations. The most differentially detected variants were cloned into new lentiviral TCR constructs based on pRRLSIN, as described in the section “Generation and use of lentiviral constructs for TCRs”.

### Discovery of new TCRs by stimulation of primary T cells by MHC I-presented peptides

To enrich T cells expressing TCRs from the centrally tolerated human TCR repertoire that could recognize HLA-A2-presented KRAS_G12V_ epitope, we isolated primary CD8^+^ T cells from HLA-A2^+^, healthy human donor PBMCs (**Fig. S4a**). From the remaining PBMCs, we generated mature DCs through a differentiation cocktail consisting of GM-CSF and IL-4 (Peprotech) for 1 d followed by TNFα, IL-1β, IL-6 (Peprotech), and prostaglandin E2 (MP Biochemicals) for 1 d. The resulting DCs acted as APCs: they were pulsed for 4 h with synthetic KRAS_G12V_ peptides (Elim Biopharm), with or without PTMs incorporated, and stimulated the CD8^+^ T cells with the resulting peptide-presenting DCs. After this initial stimulation, we continued to expand the T cells with a combination of IL-2 (Prometheus; obtained via University of Washington Pharmacy, DOH license #DRRS.FX.60550751), IL-7, and IL-15 (Peprotech) with medium replenishment every 2–3 d. These T cells were brought through multiple cycles of this stimulation. In subsequent stimulations, peptide presentation was done by peptide-pulsing PBMCs instead of mature DCs. At the end of the final round of stimulation, we stained T cells with HLA-A2 tetramers presenting KRAS_G12V_ 5-14 (with or without PTMs) and conjugated to a fluorophore, with a different fluorophore per PTM/unmodified peptide. These cells were sorted into separate populations based on tetramer binding patterns.

From the sorted T cells, we quantitated TCR clonotypes using the ImmunoSEQ assay (Adaptive Biotech) and performed single cell RNA-seq using the Chromium Single Cell V(D)J Enrichment Kit for T cells (10X Genomics) to identify paired TCRa/TCRb sequences. Using Cellranger 7.0.0 followed by Loupe VDJ Browser 4.0, for each population, we selected the most highly enriched sequences for cloning of TCR lentiviral constructs.

### Coculture with engineered TCR-T cells and HLA-A2^+^ tumor cells

HLA-A2^+^ tumor cell lines were lentivirally transduced to express either Nuclight-Red or Nuclight-Green nuclear-localizing fluorescent protein (Sartorius) and sorted for their expression. For co-culture with TCR-T cells, we plated the tumor cell lines on 48-well plates at a density of 10,000 cells/well one day before adding T cells to allow for tumor cell adherence. T cells, generated as described above, were added to these wells at a CD8^+^ T cell:tumor cell ratio of 4:1 unless stated otherwise, with an initial dose of 12.5 IU/ml IL-2. Cocultures were imaged every two hours at 10X magnification using an Incucyte S3 machine (Sartorius).

### *In vitro* peptide assays

To determine functional avidity of our engineered TCRs against peptides, we used either Jurkat CD8^+^ Nur77-GFP reporter cells or primary human HLA-A2^+^ T cells and transduced them with one of our TCRs. Cells were incubated with *TAP*-deficient T2 cells and synthetic peptide at set concentrations, in the presence of GolgiStop and GolgiPlug protein transport inhibitors (BD Biosciences) at 1:1000 dilution each for 16 h at 37°C. Cells were incubated by themselves in the presence of the same protein transport inhibitors in addition to synthetic peptides at set concentrations for 4 h at 37°C. At the end of incubation, live CD8^+^ cells were measured for expression of IFNγ (primary CD8^+^ T cells) or Nur77-GFP (Jurkat CD8^+^ Nur77-GFP reporter cells) by flow cytometry. CD8^+^ T cells were stained with eFluor 780 Fixable Viability Dye (Invitrogen) followed by CD8:BV421 (Biolegend), fixed and permeabilized using the BD Cytofix/Cytoperm Kit (BD Biosciences), and stained with IFNγ:PE-Cy7 antibody (Biolegend). Nur77-GFP Jurkat cells were stained with CD8:BV421 (Biolegend) followed by propidium iodide. Flow cytometry analysis was performed using FlowJo v9 (BD Biosciences).

To measure the stability of peptide presentation on HLA-A2, we incubated T2 cells with individual peptides of interest for 16 h, then measured live cells for surface expression of HLA-A2 by flow cytometry, staining with HLA-A2:FITC antibody (Biolegend) and propidium iodide.

### CRISPR knockout screen and validation of genes involved in methylation processes

Genes were selected using the PANTHER database (*63*) which met the criteria of containing “methyl” in the gene ontology description. The total number of genes selected was 605. A total of 1,813 sgRNA sequences for the genes of interest were collected from existing databases (*64–66*). 30 control gRNA sequences were added to the library for a total of 1,843 gRNAs (**Table S1**). The list of sgRNAs was ordered as a custom oligonucleotide pool from Integrated DNA Technologies.

For the creation of the CRISPR knockout library plasmid, we performed cloning following Canver *et al.*, 2018 (*67*). An oligo pool was cloned into lentiCRISPRv2 (Addgene #52961) which was digested using Esp3I-HF (New England Biolabs). (2.5 μg) was transfected into HEK293T cells using X- tremeGENE™ 9 (Roche) with psPAX2 (1.625 μg) and pCMV-VSV-G (0.875 μg) plasmids (Addgene #12260, #8454). Viral supernatant was collected and filtered using a 0.45 μm PVDF filter 48 hours after transfection. The plasmid-containing viral supernatant was then transduced into the DAN-G cell line. To select for the cells which were successfully transduced, the cells underwent puromycin selection for 7 days at 1 μg/ml concentration with medium changes every 2-3 days. The library-containing DAN-G cells were co-cultured with the KRAS-targeting TCR-T cells for 7 days. Following co-culture, the DAN-G cells were rinsed with PBS and pelleted for genomic DNA extraction using QIAGEN DNeasy Blood & Tissue kit. The sgRNA region of the genomic DNA was PCR amplified following the ‘sgRNA/shRNA/ORF PCR for Illumina Sequencing’ protocol via the Broad Institute Genetic Perturbation Platform and assessed for fragment length distribution on TapeStation D1000 (Agilent). Upon verification of fragment length, the samples were pooled and sequenced on an Illumina MiSeq machine.

Raw NGS reads were adapter-trimmed with Cutadapt v2.9, followed by alignment to the reference library with Bowtie v1.3. Comparisons between different conditions were performed using the MAGeCK pipeline, using RRA and MLE algorithms for calculating statistical significance of results. MAGeCK also predicts gene essentiality by evaluating the ranked list of sgRNAs (*68*).

To verify hits from our CRISPR library screen, we transduced DAN-G cells with lentivirus inducing single knockouts and puromycin resistance. After 7 days of puromycin selection at 1 μg/ml, the DAN-G single knockout cells was co-cultured with TCR-T cells at a 4:1 T cell:tumor cell ratio and imaged every 4 hours at 10X magnification using an Incucyte S3 machine (Sartorius).

To verify knockout, genomic DNA was extracted from single knockout DAN-G cells, and the gRNA region was PCR amplified using Q5 High-Fidelity DNA Polymerase (New England Biolabs), yielding amplicon lengths of between 500-800 base pairs. The PCR-amplified samples were sequenced using the “Premium PCR Sequencing” service from Primordium Labs. The resulting sequence files were aligned with the reference genome using BWA-MEM to screen for the common mutation sites to confirm the knockout. To verify the methylation status of HLA-A2-presented KRAS_G12V_ epitope in the *SUPT6H* knockout DAN-G cells and non-knocked-out DAN-G cells, we expanded these cells and extracted HLA- A2-presented peptide using the ARTEMIS protocol as previously described.

To compare HLA-A2 expression of *SUPT6H* knockout vs. non-knocked-out cells, the cells were stained with HLA-A2:PE antibody (Biolegend) and propidium iodide and analyzed via flow cytometry. To compare KRAS expression, protein lysate was extracted from tumor cell lines as described above and used as input for Western blots with anti-KRAS antibody (12063-1-AP; Thermo Fisher).

## Acknowledgements

We would like to thank the Fred Hutch Genomics, Bioinformatics, Proteomics/Metabolomics, Flow Cytometry, and Cellular Imaging Shared Resources for their respective roles in helping to run key experiments. We also thank members of the Chapuis Lab and the Strong Lab at Fred Hutch for their help setting up ARTEMIS for use in this study.

## Author contributions

Conceptualization: JWL, RP, TM, PB, AGC, TMS, PDG

Methodology: JWL, EYC, RP, MEC, TM, YS, PB, TMS, PDG

Investigation: JWL, EYC, TH, RP, MEC, TM, SM, CLM, LNJ, PRG, PC, PB, TMS

Visualization: JWL, EYC, TMS

Funding acquisition: RP, TM, AGC, PB, TMS, PDG

Project administration: AGC, TMS, PDG

Supervision: PB, TMS, PDG

Writing – original draft: JWL, EYC

Writing – review & editing: JWL, EYC, TH, RP, MEC, TM, LNJ, PRG, PC, YS, AGC, TMS, PDG

## Data and materials availability

Code for structural modeling and TCR sequence data processing from TCR mutagenesis are available at https://github.com/jihoonlee0/A2_KRAS-G12V.

## Supplementary materials

**Fig. S1:**
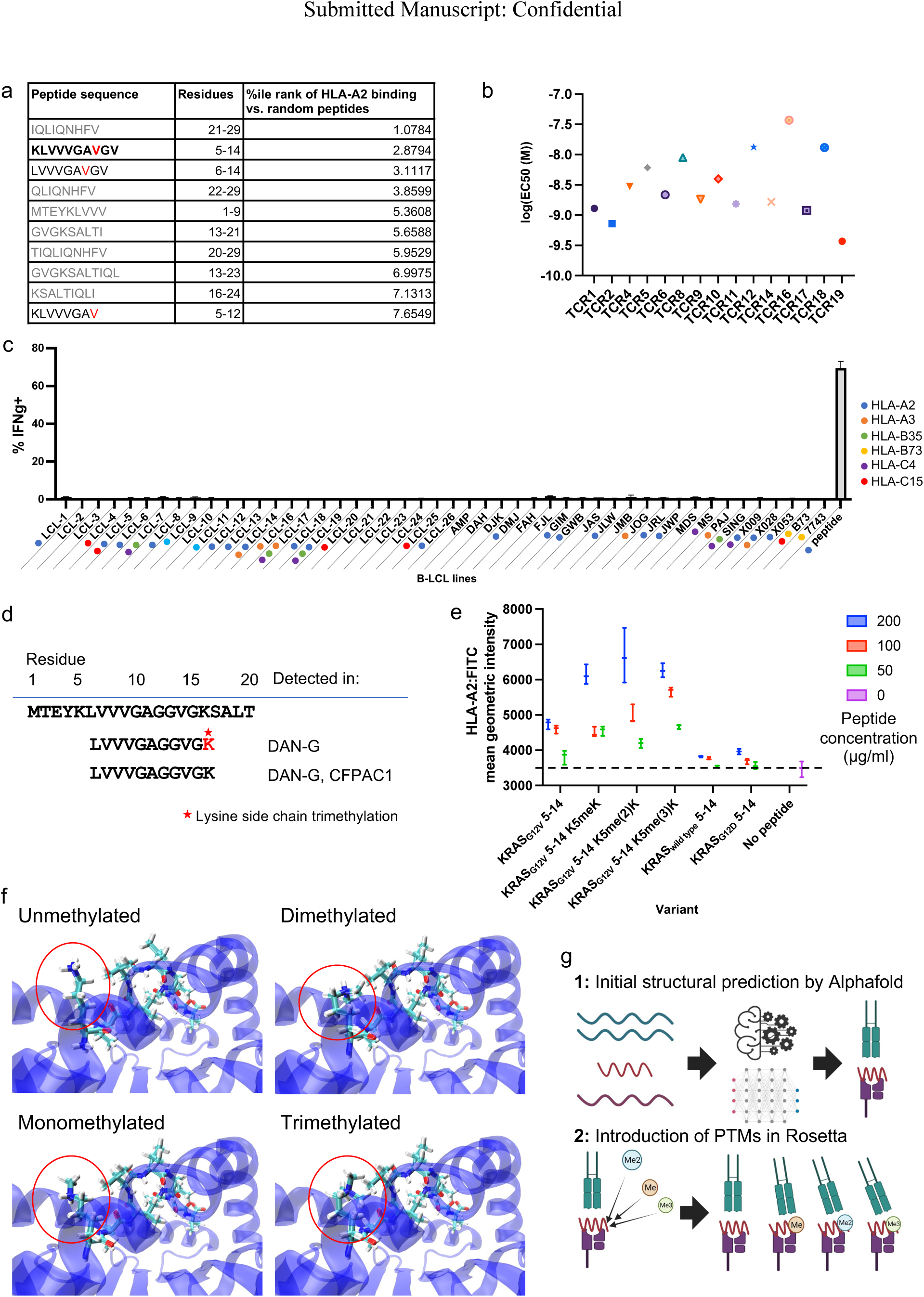
HLA-A2-presented KRAS_G12V_ epitope includes a lysine residue that can be stably presented with potential side chain methylation. (a) NetMHCPan 4.1 summary of the top 10 peptide sequences that are most likely to be presented by HLA-A2 from the first 30 amino acids of KRAS_G12V_. Peptides containing the G12V mutation (including mutations in red text) are in black text; others are in gray. (b) Log_10_(EC_50_ (M)) comparison of recognition of synthetic KRAS_G12V_ 5-14 peptide (KLVVVGAVGV) presented by HLA-A2 by an initially generated set of MHC class I-restricted TCRs isolated to be specific for this epitope. Lower values indicate higher functional avidity. (c) % IFNγ^+^ T cells, transduced with TCR_2_, after exposure to B-LCL cell lines expressing different Class I HLA alleles found on CFPAC1 cells without KRAS_G12V_ 5-14 peptide loaded. Cell lines are color coded by allele. (d) MS results of RAS family peptide fragments detected after tryptic digestion of immunoprecipitated whole KRAS protein from cell lysate of DAN-G and CFPAC1 pancreatic adenocarcinoma cell lines. Lysine side chain trimethylation is indicated in red with an asterisk. (e) Reconstitution of properly folded HLA-A2 surface expression on T2 cells after exogeneous loading by wild type or G12V/D mutant KRAS 5-14 peptides with or without lysine-5 side chain methylations. (f) Additional views of structural models of HLA-A2-presented KRAS_G12V_ 5-14 peptide including possible lysine-5 side chain methylation, supplementing Fig. 1e by highlighting the protrusion of the lysine-5 side chain (circled in red) out of the HLA-A2 groove. (g) Overview of structural prediction of a TCR-pMHC complex using a combined Alphafold-Rosetta pipeline. Alphafold allows for deep learning-based prediction of proteins and protein complexes, which we applied to predict TCR-pMHC complex structure. However, this does not yet contain PTMs, which Alphafold cannot incorporate reliably. Thus, we used the resulting structure as input into Rosetta, which used physics-based energy functions to introduce PTMs and reoptimize the structure to accommodate them.

**Fig. S2:**
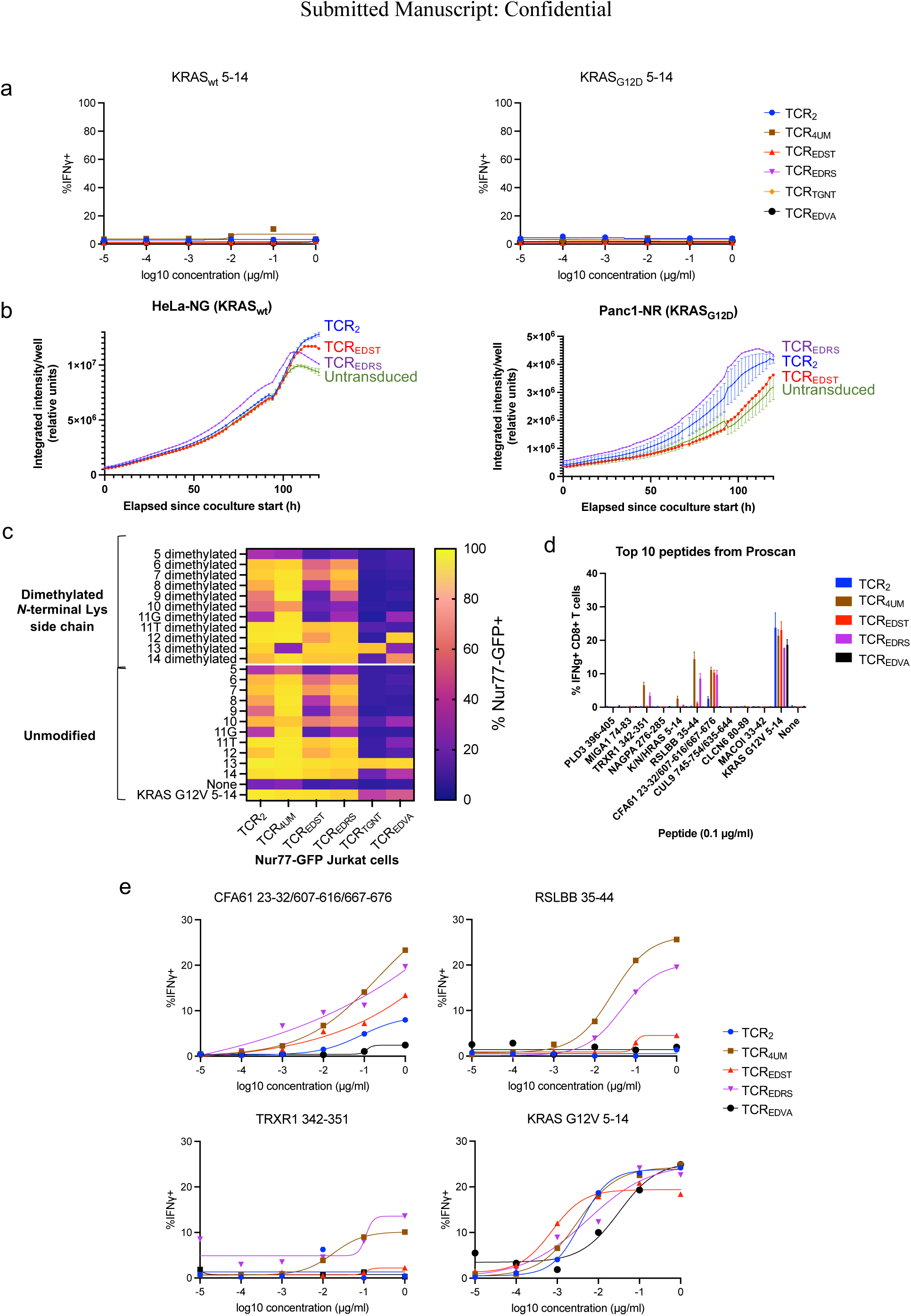
TCRs developed from a mutagenesis library to recognize post-translationally modified epitopes of HLA-A2-presented KRAS_G12V_ are G12V mutation-specific. (a) Dose response curves of CD8^+^ T cells expressing selected TCRs generated from the HLA-A2 KRAS_G12V_ TCR mutagenesis library after exposure to wild type or G12D KRAS 5-14 peptides presented by HLA-A2. (b) Failure of CD8^+^ T cell killing and progressive growth of human HLA-A2^+^ tumor cell lines that express wild type KRAS (HeLa cervical carcinoma cell line; left) or KRAS_G12D_ (Panc1 pancreatic adenocarcinoma; right) and no KRAS_G12V_. Cocultures of T cells and tumor cells were recorded by Incucyte imaging. Tumor cells expressed nuclear fluorescent protein. NR: red fluorescent protein; NG: green fluorescent protein. (c) Alanine scan of KRAS_G12V_ 5-14 epitope tested against TCRs selected from the mutagenesis library. Each cell indicates the resulting expression of Nur77-GFP upon replacement of a KRAS_G12V_ residue with alanine (or in the case of alanine-11, replacement with either glycine or threonine) and with or without lysine-5 side chain dimethylation. (d) IFNγ secretion by CD8^+^ T cells expressing mutagenized TCRs after exposure to synthetic peptides from the human proteome fitting the K-x-x-V-V-x-A-x-x-x tolerance pattern identified from the alanine scan shown in (c), with a concentration of 0.1 μg/ml for each peptide. (e) Dose response curves of CD8^+^ T cells expressing mutagenized TCRs after exposure to the potential human off-target peptides demonstrated above to generate the three largest responses at 0.1 μg/ml: CFA61_23-32/607-616/667-676_, RSLBB_35-44_, and TRXR1_342-351_, compared to the KRAS_G12V_ epitope.

**Fig. S3:**
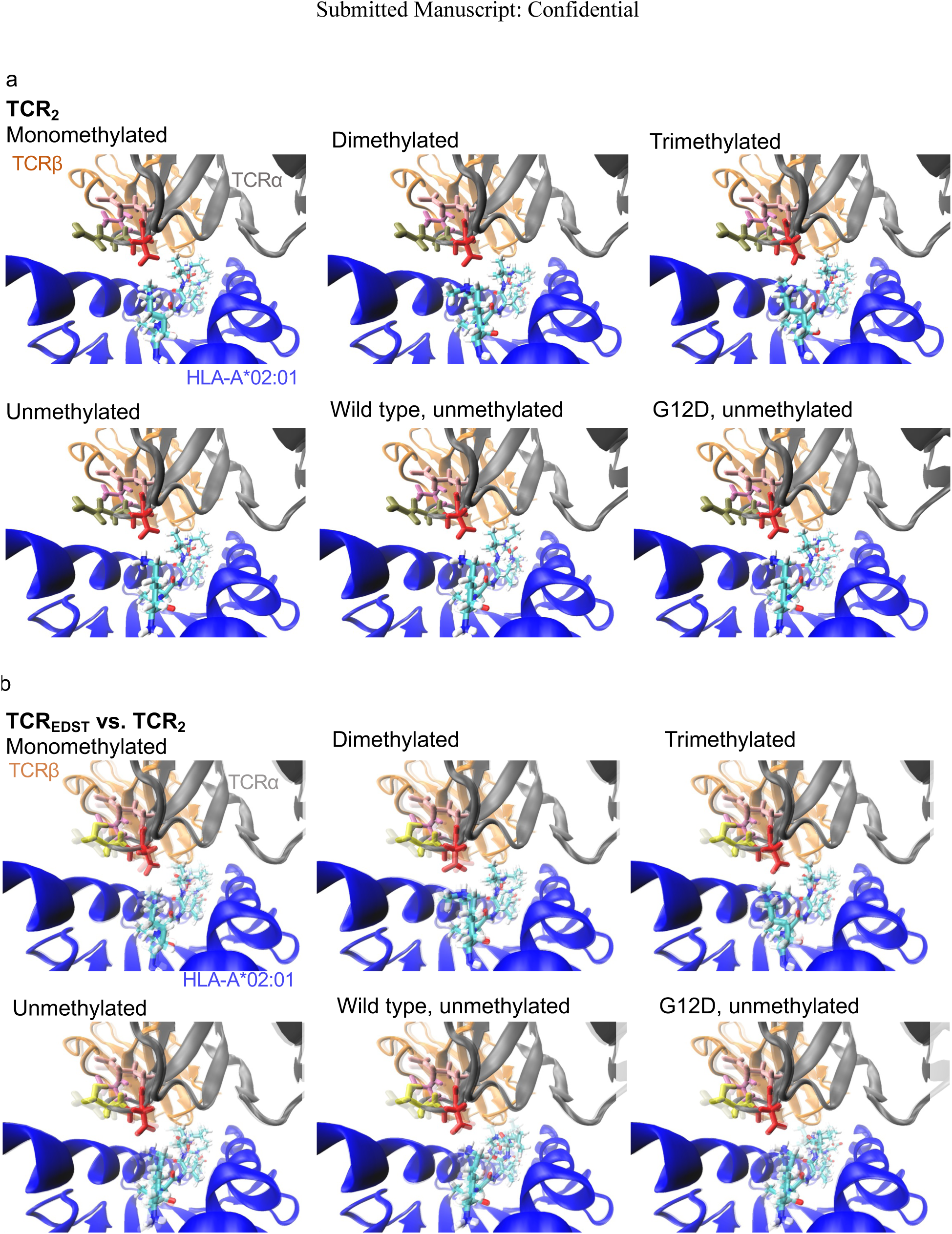
TCR-pMHC structural models by Alphafold-Rosetta show CDR3α core interactions with N-terminal lysine side chain of HLA-A2-presented KRAS_G12V_ epitope. Structural predictions of unmethylated and methylated variants of KRAS_G12V_ 5-14, as well as unmethylated wild type or G12D peptide, presented by HLA-A2 to interface with (a) TCR_2_ and (b) mutagenized TCR_EDST_ TCRs. The CDR3α ‘core’ sequence is shown as individual amino acids within the TCRs. In (b), the TCR_2_-pMHC interfaces from (a) are overlaid with transparency to facilitate visual comparison. With TCR_EDST_, residues in the core sequence of CDR3α and the lysine-5 side chain of each variant of KRAS_G12V_ epitope appear closer to each other than with TCR_2_.

**Fig. S4:**
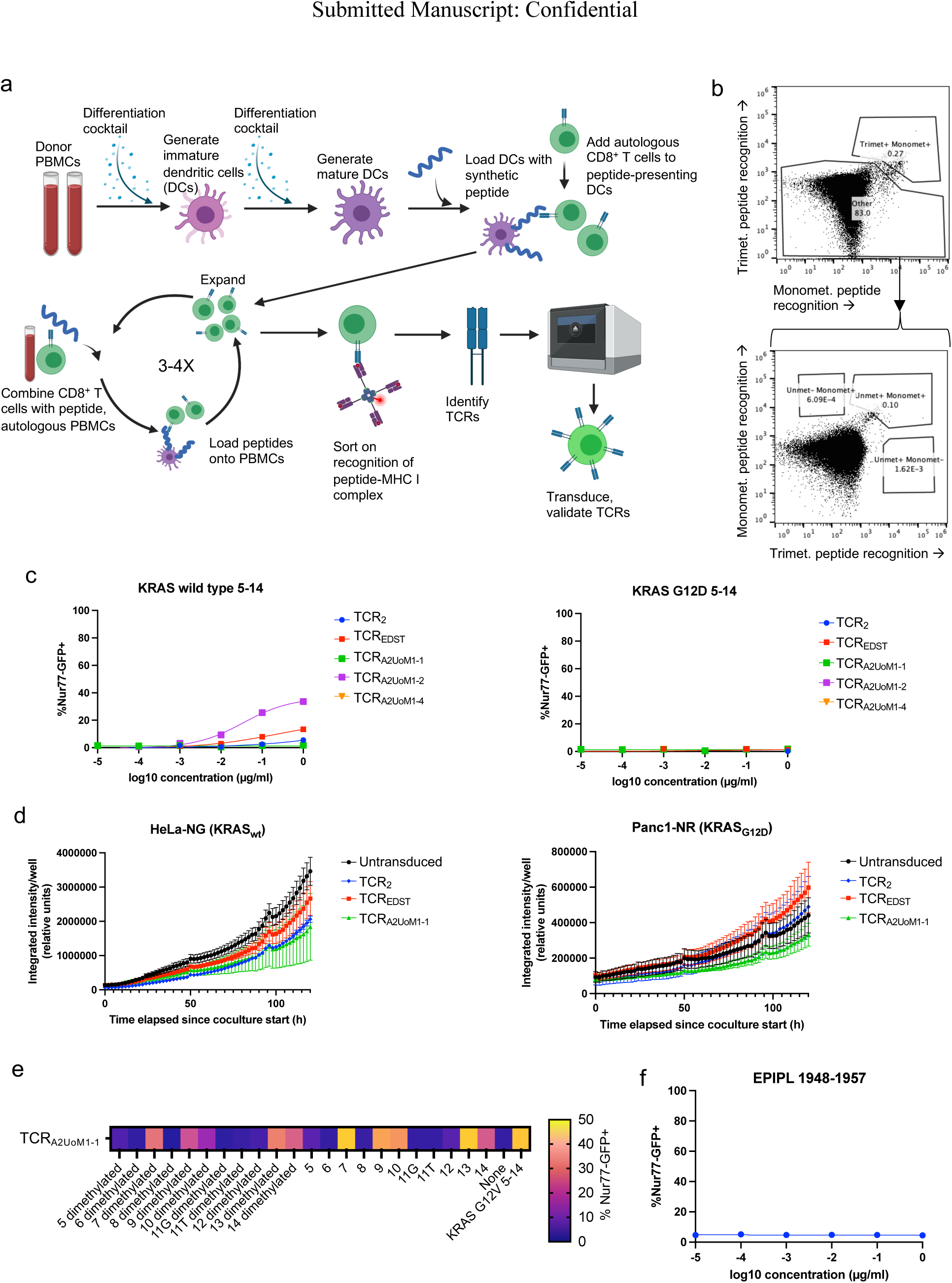
TCRs identified from primary human T cells specifically target KRAS_G12V_ epitope presented by HLA-A2. (a) Experimental workflow to enrich and characterize T cell clones with TCRs recognizing a defined peptide epitope. (b) Tetramer binding patterns of primary human T cells stimulated with unmethylated and methylated KRAS_G12V_ peptide presented by HLA-A2. Plots also indicate gates used to sort cells with different affinities for unmethylated and methylated epitopes. (c) Dose response curves of TCRs against wild type and KRAS_G12D_ 5-14 peptides presented by HLA-A2. (d) Tumor cell growth during coculture of CD8^+^ T cells expressing selected TCRs with HLA-A2^+^, wild type KRAS HeLa and KRAS_G12D_ Panc1 pancreatic adenocarcinoma cell lines. Cocultures of T cells and tumor cells were recorded by Incucyte imaging. Tumor cells expressed nuclear fluorescent protein. NR: red fluorescent protein; NG: green fluorescent protein. (e) Alanine scan of KRAS_G12V_ 5-14 epitope tested against TCR_A2UoM1-1_. Each box indicates the resulting expression of Nur77-GFP upon replacement of a KRAS_G12V_ residue with alanine (or in the case of alanine-11, replacement with either glycine or threonine). (f) Dose response curve of CD8^+^ Nur77-GFP Jurkat cells expressing TCR_A2UoM1-1_ after exposure to EPIPL_1948-1957_, the potential human off-target peptide as determined from alanine scan results.

**Fig. S5:**
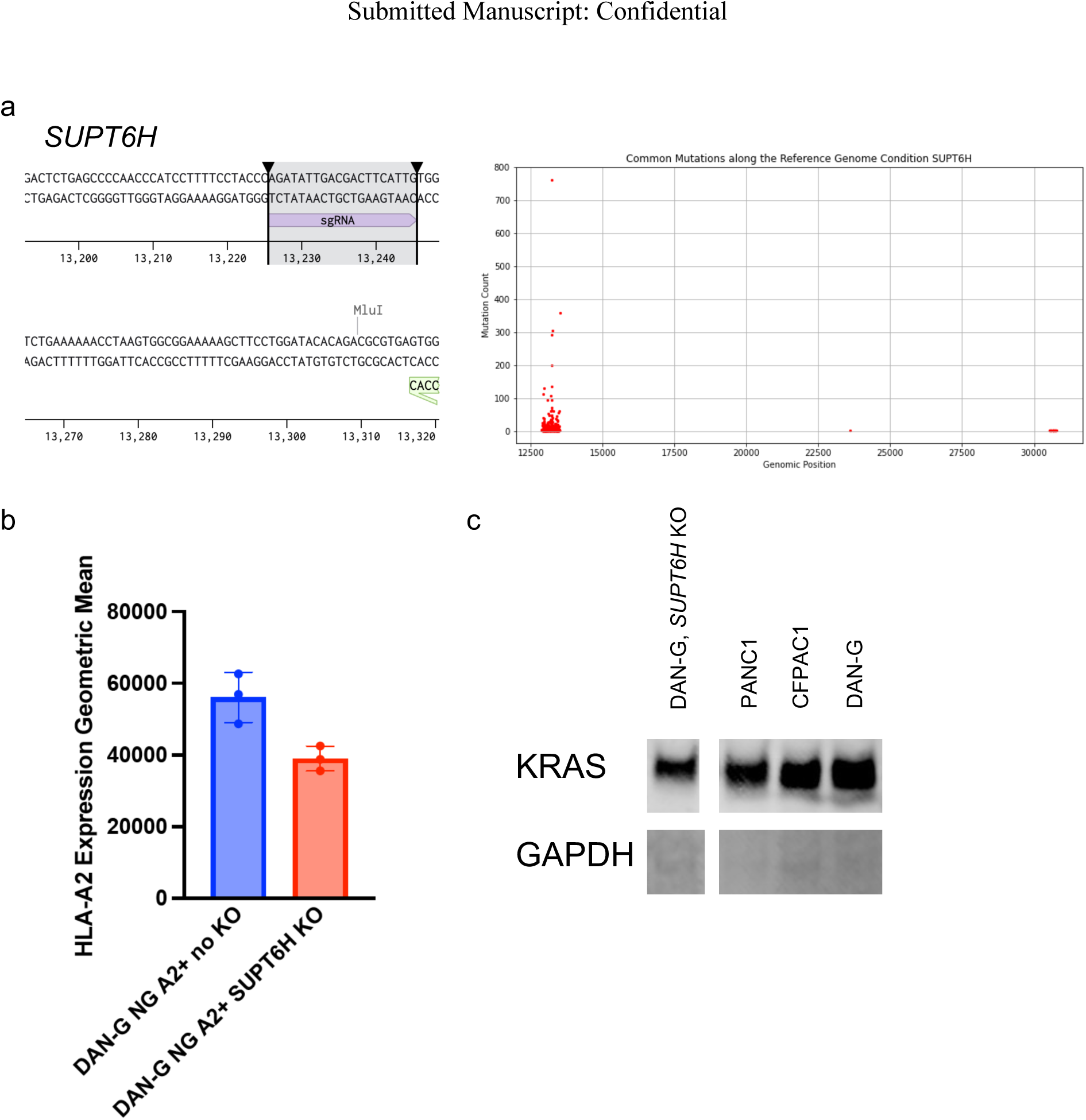
Further characterization of CRISPR knockout of *SUPT6H*. (a) Location of the gRNA binding region in the *SUPT6H* gene and summary of the genomic position of the commonly occurring mutations detected from DAN-G cells transduced with lentiCRISPRv2 with gRNA targeting *SUPT6H*. Red dots indicate a mutation detected from long-read sequencing. (b) Surface expression of HLA-A2 with and without *SUPT6H* knockout in DAN-G cells. (c) Western blot of total expressed KRAS protein with and without *SUPT6H* knockout in DAN-G cells. KRAS expression in other pancreatic adenocarcinoma cell lines Panc1 and CFPAC1 are shown for additional comparison. GAPDH protein expression was used as a control.

**Table S1:** List of gRNAs for CRISPR knockout screen of methylation-related genes.

